# Transposase N-terminal phosphorylation and asymmetric transposon ends inhibit *piggyBac* transposition in mammalian cells

**DOI:** 10.1101/2022.09.26.509586

**Authors:** Wentian Luo, Alison B. Hickman, Pavol Genzor, Rodolfo Ghirlando, Christopher M. Furman, Anna Menshikh, Astrid Haase, Fred Dyda, Matthew H. Wilson

## Abstract

Mechanistic regulation of DNA transposon systems in mammalian cells remains poorly understood. Using modeling, biochemical, and cell-based assays, we sought to extend the recent cryoEM structural insight into the *piggyBac* transpososome to evaluate the previously unexplained role of the transposase N-terminus, the need for asymmetric transposon ends, and the complexity of transposase tetramer formation for transposition in mammalian cells. We found that N-terminal phosphorylation by casein kinase II inhibits transposase-DNA interaction and designed deletion of this phosphorylated domain releases inhibition thereby enhancing activity. We also found that the N-terminal domain promotes transposase dimerization in the absence of transposon DNA. N-terminal deletion enables transposition of symmetric transposon ends that was previously not achievable with *piggyBac*. The complex transposase tetramer needed for transposition of asymmetric transposon ends can be overcome via appending a second transposase C-terminal domain in combination with symmetric transposon ends overcoming the negative regulation by asymmetric ends. Our results demonstrate that N-terminal transposase phosphorylation and the requirement for asymmetric transposon ends both negatively regulate *piggyBac* transposons in mammalian cells. These novel insights into mechanism and structure of the *piggyBac* transposase expand its potential use for genomic applications.

## Introduction

Genomes are not static partly due to transposable elements that can move discrete pieces of DNA (called transposons) from one place to another or coordinate the generation of another transposon copy at a new location. Although transposons have been shown to contribute to adaptation, they are usually considered neutral or detrimental to host species. For example, although integration into a new site brings the potential for adaptive rewiring of regulatory pathways, it can alternatively result in genotoxicity. Thus, many transposons have evolutionarily drifted away from maximal activity, increasing their chances of co-existing within their hosts while maintaining some ability to move and avoid host-imposed regulation. Understanding of the mechanistic regulation of DNA transposons in mammalian cells remains incompletely understood.

The existence of transposons presents opportunities for their use in genome engineering that can complement other DNA-targeting systems such TALE nucleases or CRISPR-Cas based tools (1,2). The most widely used transposons in higher organisms are *Sleeping Beauty* (SB) (3,4), a resurrected mariner transposon originally identified as inactive in fish, and *piggyBac* (PB) (5-7), an active transposon isolated from the cabbage looper moth. PB exhibits unique properties compared to other DNA transposons including specificity for insertion at the tetranucleotide sequence TTAA and the advantageous ability to couple its genomic excision with seamless repair in the host cell (5,6). PB has been used for a wide range of biotechnology applications including generation of transgenic animals, functional genomics, cancer gene discovery, and cell and gene therapy applications (7-13).

Mechanistic regulation of PB can occur at the host cellular level or within the PB transpososome itself (transposase interaction with transposon DNA). At the host level, PB has been shown to interact with DNA-dependent protein kinase for paired-end complex formation (14). PB also interacts with bromodomain containing proteins (i.e. BRD4) which appear to bias PB integration towards known sites of genomic DNA interaction with BRD4 (15). Within PB itself, analysis has revealed the nuclear localization site (16), the catalytic core residues (17), DNA binding sites (18), amino acids enabling transposon excision without integration (19), and the lack of the need for DNA synthesis for transposition (20). Hyperactive PB has also been generated via random transposase mutagenesis (21-23), peptide addition (24), or manipulating transposon ends (25). Nonetheless, there remain unanswered questions regarding PB-mediated transposition in mammalian cells.

The wild-type (WT) PB transposon consists of an ORF encoding its transposase flanked by dissimilar transposon ends that contain several short DNA motifs arranged asymmetrically on its Left End (LE) and Right End (RE) (**Fig. 1a**); this asymmetry is crucial for transposition activity in cells (26). The recent cryo-EM structures of the PB transposase bound to symmetric transposon hairpin ends and in a strand transfer complex with TTAA-containing target DNA provided not only insight into its mechanism of excision and integration, but also suggested the need for an active assembly (or transpososome) containing four PB protomers as the rationale for the end asymmetry requirement (26). This model was derived due to the fact that structure was obtained with symmetric transposon ends though activity of transposition of symmetric transposon ends was not achievable (26). These first PB transpososome structures also left an absence of density for the first 116 N-terminal transposase residues, leaving the function of this domain unidentified (26). A recent investigation of the N-terminal 100 amino acids (AAs) demonstrated a loss of activity with deletion but without mechanistic explanation (27).

**Fig. 1.**
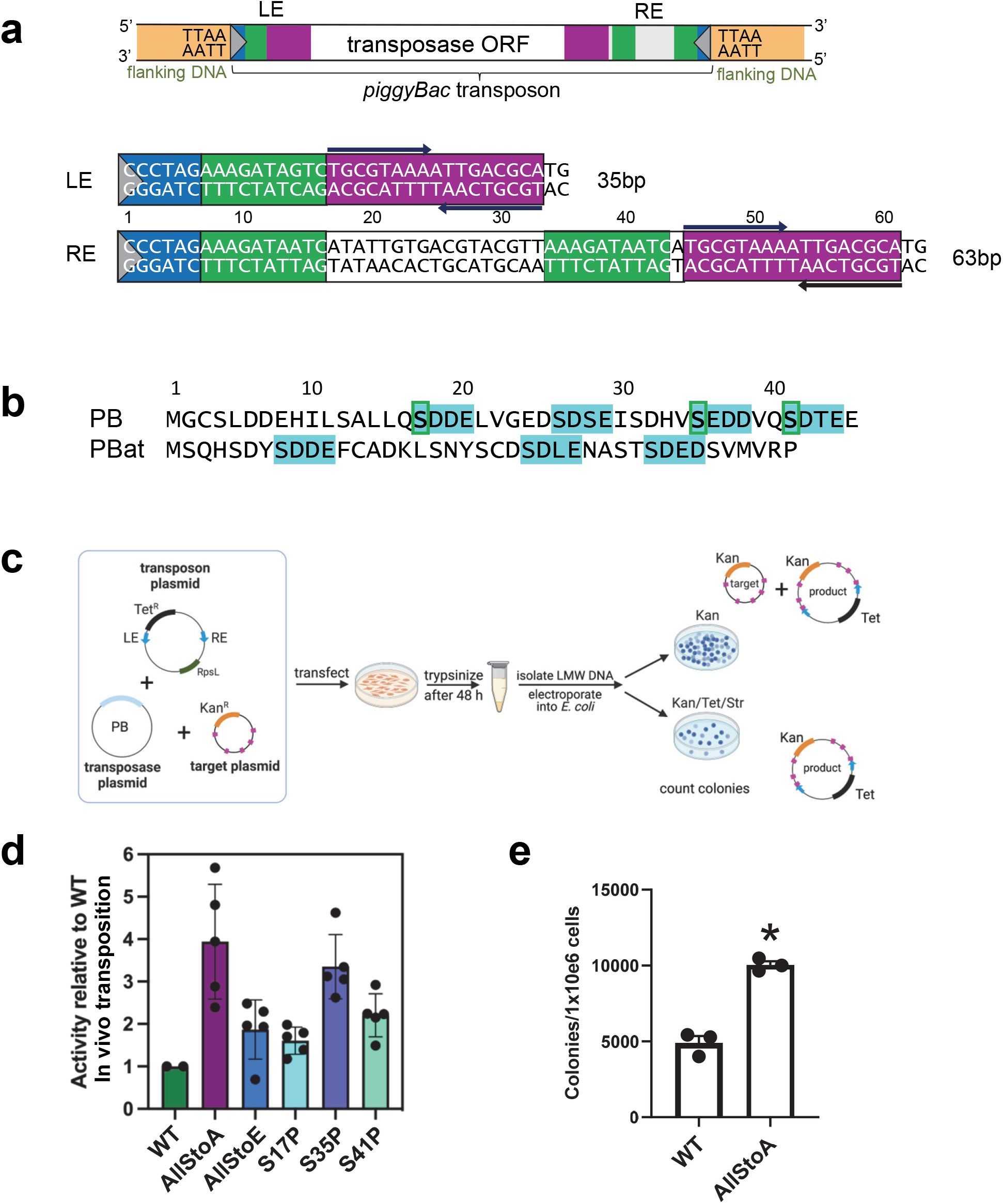
*piggyBac* transposon organization, and identification of transposase N-terminal CKII dependent phosphorylation that inhibits activity in human cells. **a**. Schematic of PB transposon flanked by TTAA, and sequence and organization of the LE and RE TIRs. **b**. Alignment of the N-terminus of PB and PBat demonstrating CKII phosphorylation sites (highlighted). CKII sites within PB found to be phosphorylated when expressed in human cells are marked with a green box. **c**. Schematic of inter-plasmid in cell transposition assay. Isolated episomal DNA post-transfection was electroporated into bacteria which were plated on medium containing kanamycin to measure the total recovery of recipient plasmids or on kanamycin/tetracyline/streptomycin plates. Tet and Kan allow selection for the transposition product, and streptomycin selects against donor plasmids which contain the streptomycin sensitivity gene *rpsL*. **d**. In cell transposition activity comparing phosphorylation site mutations to WT using a inter-plasmid transposition assay in human cells. **e**. Colony count (integration) assay of a neomycin resistant transposon comparing AllStoA PB to WT PB in human cells. N=3±SEM; *, p<0.05 using student’s T test.

We undertook the current investigation to advance beyond the recent cryoEM insight into the PB transpososome to further understand the previously unexplained functional role of the transposase N-terminus, the need for asymmetric transposon ends, and the complexity of the transposase tetramer formation within the transpososome for transposition in mammalian cells. We also sought to simplify the PB transpososome complex to advance its future capabilities for genome engineering.

## Materials and Methods

### Plasmid constructs

pCMV-HAPB and pTpB with LE-RE and LE-LE TIRs have been described previously (26,28). pCMV-hyPB has been described previously(22). pCMV-Δ74PB and pCMV-Δ74m7pB were generated by deleting N-terminal amino acids 1-74 using PCR while retaining an initiation methoinine. pCMV-PB-2CD and pCMV-hyPB-2CD were generated by adding amino acids 542-594 of PB to the end of C-terminus in tandem. pCMV-Δ74PB-2CD and pCMV-Δ74hyPB-2CD were generated by deleting N-terminal amino acids 1-74 and adding amino acids 469-521 of PB to the end of C-terminus. pT-mAppleT2Apuro was generated by PCR amplifying the T2A peptide and puromycin resistance gene from PB-CMV-MCS-GreenPuro (System Biosciences, Cat# PB513B-1) and cloning into pT-mApple (Vectorbuilder). pT-mAppleT2Apuro-β-geo was generated by cloning the splice acceptor-β-geo fragment from PB-SB-SA-βgeo(29) into pT-mAppleT2Apuro. pT-mAppleT2Apuro(15.1kb) was generated by cloning a PacI/BamHI restriction enzyme fragment from pAdEasy-1 (Addgene, Plasmid#16400) into pT-mApple. PB-SRT-Puro LE-RE and PB-SRT-Puro LE-LE were generated by replacing full length TIRs of PB-SRT-Puro (30) using shorter LE and/or RE TIRs. Standard molecular biology techniques were used, and all constructs were confirmed with DNA sequencing.

For the plasmid-to-plasmid in cell assay, pFV4a-PB was synthesized by GenScript, and was derived from the Helraiser (HR) transposase expression plasmid, pFV4aRH (31), by exchanging the HR ORF for that of human codon-optimized PB. pFV4a-PBAllStoA, pFV4a-PBAllStoE, pFV4a-PBS17P, pFV4a-PBS35P, and pFV4a-PBS41P were generated by ligation of the appropriate gBlock (IDT) between the SpeI and BmtI sites of pFV4a-PB. The donor plasmid, pTet-pBac-LE35-RE63, was synthesized by GenScript and was generated by replacing the 12- and 12-RSSs of pTet-RSS (32) with PB LE35 and RE63. Both pTet-RSS and the target plasmid, pHSG298, were kind gifts of the David Schatz lab. pD2610-MBP-PB has been described previously (21). pD2610-MBP-PB1-539, pD2610-MBP-TEVΔ74PB, and, pD2610-MBP-TEVΔ74PB1-539 were synthesized by Twist Bioscience; for the TEVΔ74 constructs, the sequence ENLYFQG was inserted between amino acids G74 and S75.

### Protein purification

PB1-539, TEVΔ74PB, and TEVΔ74PB1-539 were expressed in Expi293F cells (Thermo Fisher) and purified as previously described (26).

### Mass spectrometry analysis

Phosphorylation of purified PB1-539 was assessed by peptide digestion and micro-capillary LC/MS/MS (Taplin Mass Spectrometry Facility, Harvard Medical School). Phosphorylation sites were assigned based on their Modscore values (33). The level of phosphorylation was estimated based on the peak intensities for the same peptide with and without phosphate.

### Sedimentation velocity analytical ultracentrifugation (SV-AUC)

Purified PB1-558 in 500 mM NaCl, 20 mM HEPES.NaOH pH 7.6 and 0.5 mM TCEP, and PB74-539 in 500 mM NaCl, 25 mM TRIS.HCl pH 7.4 and 0.5 mM TCEP were prepared at ∼5.5 µM. Sedimentation velocity data were collected at 50,000 rpm and 20°C on a Beckman Coulter ProteomeLab XL-I analytical ultracentrifuge following standard protocols (34). 2-channel, 12 mm centerpiece cells were used, and sedimentation was monitored with the absorbance (280 nm) and Rayleigh interference (655 nm) detection systems. A continuous c(s) distribution of Lamm equation solutions, as implemented in SEDFIT (35), modeled the sedimentation data. SEDNTERP(36) provided the solution densities ρ, solution viscosities η, and protein partial specific volumes required for the analysis.

### AlphaFold-Multimer

AlphaFold-Multimer (5×5) (37) was run on the NIH High Performance Computing system to generate 25 structural models for the full-length PB dimer from five different random seeds; the pLDDT scores ranged from 0.808 to 0.698. Among the ten models with the highest scores, three had regions in trans. Of these, the highest-ranking model is shown in **Fig. 2a**; two other models placed the first α-helix, residues 7-15, in trans.

**Fig. 2.**
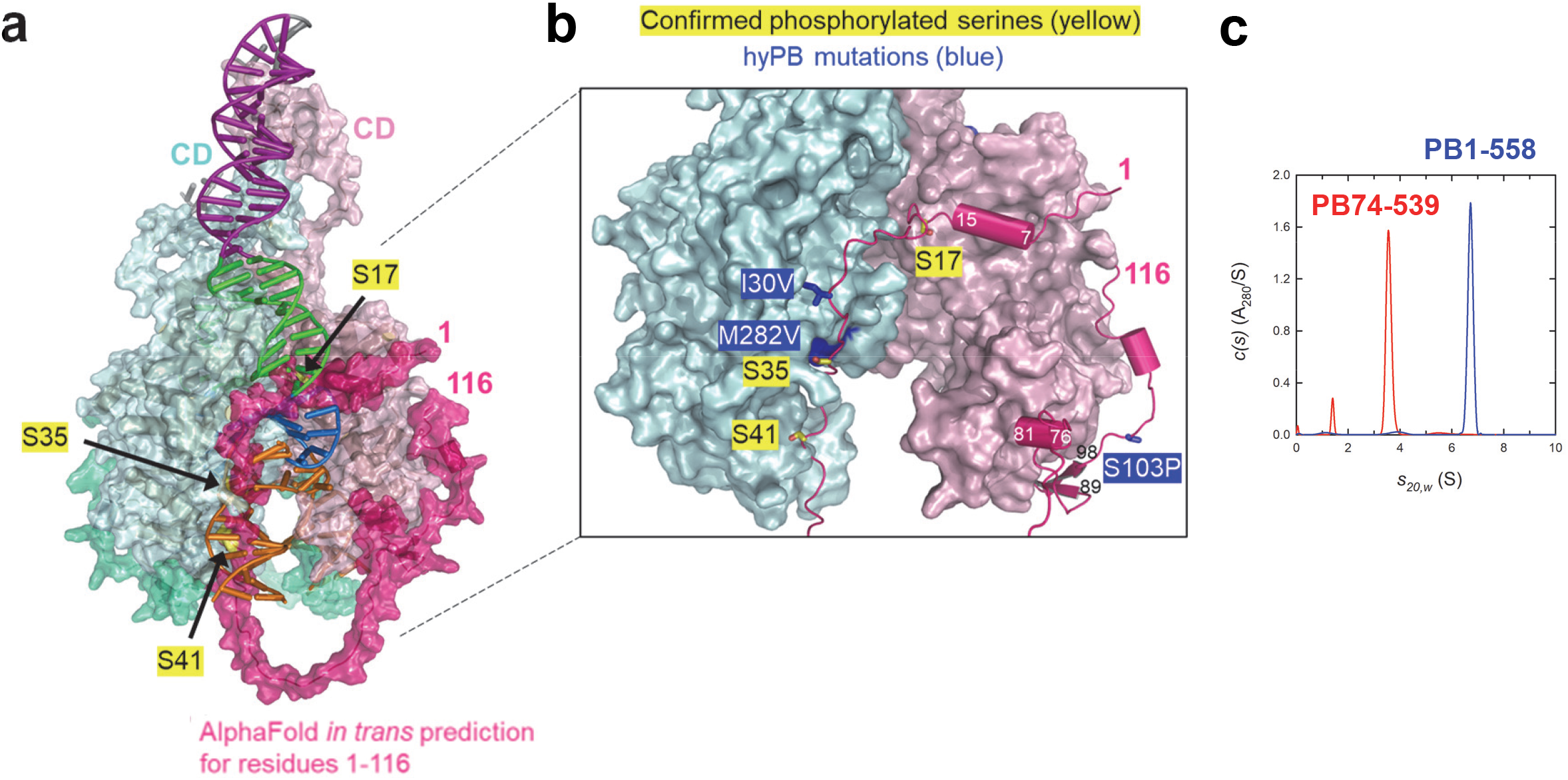
AlphaFold structural prediction for PB N-terminal region suggests multiple roles. **a** and **b**. AlphaFold-Multimer modeling of full length PB suggests that N-terminus phosphorylation inhibits DNA binding. **c**. Absorbance sedimentation c(s) profiles for 5.8 µM PB1-558 (blue) and 5.4 µM PB74-539 (red) show the presence of a dimer and monomer, respectively. Data collected for 1.4 µM PB4-539 showed a profile similar to that for the more concentrated sample.

### Cell culture and transfection

HT-1080 cells were cultured using standard procedures (38). For transfection (unless otherwise indicated), cells were seeded at a density of 300,000 cells per well in a six-well plate and transfected with 2.5 µg of total plasmid DNA, containing 1.5µg of transposon and 1 µg of transposase plasmid DNA unless otherwise indicated using Lipofectamine LTX (Invitrogen), according to manufacturer’s instructions. For varying tranposase DNA amount, cells were transfected with 1.5µg of transposon and various transposase plasmid DNA amounts with pUC plasmid DNA added to make the total DNA amount 4µg per condition. Cells were trypsinized and re-plated for functional assays 24 h later. For comparing different transposon sizes, cells were transfected with 1.0µg of transposases and transposons mAppleT2APuro (0.65µg) or mApple-β-Geo(1.1µg) or mAppleT2APuro pAdeasy(1.8µg) to keep the number of transposon plasmids equivalent between transfections. The amount of DNA was added to 2.8µg using pUC19 as remainder DNA.

The plasmid-to-plasmid assay was performed essentially as described (32). HEK 293T cells were plated at a density of 500,000 cells per well in a six-well plate and transfected the next day with 0.5 µg transposase plasmid, 1 µg transposon plasmid, and 1.5 µg target plasmid DNA using Lipofectamine 3000 (Invitrogen) according to the manufacturer’s instructions. Cells were collected 48 h later, and LMW DNA isolated using a modification of Birnboim and Doly, 1979(39). After the third ethanol-precipitation step, the pellet was redissolved in 25 µl of TE buffer (0.01 M Tris-HCl pH 8.0, 1 mM sodium EDTA) and used for electroporation of ElectroMAX DH10B cells (Invitrogen). Cells were subsequently plated on both KTS (30 µg/ml kanamycin, 12 µg/ml tetracycline, 30 µg/ml streptomycin) plates (0.4-0.8 ml) and 30 µg/ml Kan plates (1×10^4^ dilution); after 24 h, colonies were counted, and activity quantified by dividing the number of KTS colonies by the number of Kan colonies.

### Excision assay

Excision assay analysis was performed as described by previously (26,28). Plasmid DNA was recovered from transfected cells 24 h after transfection and subjected to excision PCR analysis (primers listed in **supplementary table**). PCR products were visualized using agarose gel electrophoresis. Excision bands were excised and PB transposition was confirmed via DNA sequencing as described previously (26,28).

### Colony count assay

One day after transfection, 2500 cells were replated on 10-cm dishes in growth media plus G418 (700 µg/ml) or puromycin (3µg/ml) and selected for 10 days. Dishes were then fixed, stained with methylene blue and counted as described previously (38).

### Quantitation of transposon copy number with droplet digital PCR

We used droplet digital PCR for transposon and RNAse P copy number. HT-1080 cells were transfected and selected as described above. After a minimum of 2 weeks of selection, genomic DNA was isolated as above. To reduce episomal transposon DNA, isolated DNA was treated with restriction enzyme Dpn I for which mammalian genomic DNA cleavage is blocked by overlapping CpG methylation. Ten nanograms of genomic DNA was used to amplify the neomycin resistance or RNase Ps genes (all primers, **supplementary table**). Primer/probe concentration was 900nM/250nM. Neo primers/probe and RNase P primers/probe were placed in one tube with channel 1 for Neo-FAM and channel 2 for RNAse P-Hex to reduce pipetting errors. The Neo copy number per RNAse P was directly calculated by Neo copy number divided by RNAse P copy number in 20µl reaction.

### Genome-wide sequencing library preparation

HCT116 cells were transfected with transposase plasmids pCMV-PB, hyPB, Δ74PB-2CD, Δ74hyPB-2CD and transposon plasmids PB-SRT-Puro LE-RE, PB-SRT-Puro LE-LE using lipofectamine LTX in 100mm dishes. Next day after transfection, cells were split into four 100mm dishes containing 3ug/ml of puromycin medium. After two weeks selection with puromycin, cells were collected, and RNA was prepared using an RNeasy RNA mini kit (Qiagen). Four micrograms of total RNA were used were prepared cDNAs using M-MLV reverse transcriptase, RNase H minus, point mutant which is used in cDNA synthesis with long RNA templates of more than 5kb (Promega #M3681) and primer SMART-dT18VN, (primers, **supplementary table**). cDNAs were PCR amplified with four primers located in transposon areas, SRT-PAC-F1, SRT-Seq P1, SRT-Seq P2, and SRT-Seq P3 and one primer located in Smart-dT18VN, being Smart. Amplified cDNAs were purified using PCR/Gel purification column (Macherey-Nagel #740609). 500ng of PCR amplicons were fragmented and tagged with Illumina DNA Prep (Illumina #20060060). Tagged DNA fragments were further PCR amplified using Read1-TnME, and Read2-R-5’TIR and PCR amplicons were 100-500bp size selected with 2% agarose gel. Dual index primers (NEB #E7780S) for sequencing on Illumina MiSeq and NovaSeq platforms were added by amplifying the tagmented samples using Q5 polymerase (NEB #M0544S) following NEBnext protocol (NEB #E7645S – Section 4.1). The PCR products were purified using AMPure XP beads (Beckman Coulter #A63881) using 0.9x bead volume. The quality of the final library was verified on TapeStation.

### Trimming of the raw reads

Raw reads from each sample were trimmed twice using cutadapt (40). First trimming was to remove the sequencing adapters and retain the Tn5 tagmentation sequence and transposon inverted repeat (TIR) sequence for analysis of the transposon integration features. Second trimming removed the constant TIR sequences to enable efficient alignment of the reads to the genome. Specific trimming parameters for PB are in the supplementary version of computational methods (PDF of all code) or online in the associated github page (https://github.com/HaaseLab/piggyBac_mutants).

### Sequence logo of integration preferences

The fastq file of the second read (R2) containing TIR sequence after removal of sequencing adapters was loaded into R and analyzed using the ShortRead package (41). The length of each read was trimmed to same length and reads containing perfectly matching TIR sequence at their five prime ends were selected. Nucleotide frequency per position was calculated and plotted using ggplot (42) and ggseqlogo (43) packages.

### Peak Calling

The TIR sequence was removed and reads were aligned to the human genome (Gencode GRCh38.p5.v24) in paired-end mode using Hisat aligner (44) with standard settings. The bam files were loaded into R (45) using custom function implementing Rsamtools (46), BSgenome.Hsapiens.UCSC.hg38 (47), GenomicRanges (48), GenomicAlignments (48), and data.table (49). Briefly, perfectly mapping and perfectly paired primary alignment reads were loaded into R, and each read pair converted into a single read fragment. Only unique genomic read fragments smaller than the median library size were kept for further analysis. This data was used to calculate stranded coverage that was consecutively used to find peaks using slice() function from GenomicRanges package. Any genomic position with minimum of five unique fragments coverage was identified to be a peak and used in consecutive analyses. Annotated peak list for each sample is provided with GEO (GSE201914).

## Results

### PB transposase is multiply phosphorylated in mammalian cells

One surprising aspect of the first PB transpososome structures was the absence of density for the first 116 N-terminal residues, representing 20% of the 594 amino acid transposase, indicating complete disorder in both the transposon end and strand-transfer complexes (26). Assignment of function to this segment remained elusive as observed protein/DNA interactions in the structures accounted for the available footprinting data (18) and the estimated pI of 4.3 for this protein segment suggested no role in DNA binding. A low pI for the N-terminal region of the transposase is a common property of members of the large superfamily of *piggyBac*-like elements (PBLE) although these regions cannot be aligned (50). However, we noticed that most PBLEs possess multiple casein kinase II (CKII) phosphorylation motifs, S/T-D/E-X-E/D (51), in this region (**Fig. S1**). This includes PB and *piggyBat* (PBat), two transposons in the PBLE with demonstrated activity, which have several CKII consensus sites within their transposase N-termini (**Fig. 1b**).

We used mass spectrometry to evaluate the phosphorylation status of PB expressed in HEK293F cells. The absence of basic residues in the N-terminus prevented the use of trypsin digestion, but digestion of purified PB1-539 with Asp-N protease yielded several phosphorylated peptides that could be confidently identified by LC-MS/MS (**Table S1**). Furthermore, the detection of corresponding pairs of phosphorylated vs. non-phosphorylated peptides allowed estimation of the extent of phosphorylation. The data indicated that Ser17 and Ser35 were fully phosphorylated (green boxes, **Fig. 1b**), whereas Ser41 was phosphorylated in ∼20% of spanning peptides; in this same region, Ser26 lacked detectable phosphorylation.

### N-terminal phosphorylation inhibits transposition in cells

To determine if N-terminal phosphorylation affected PB activity, we used a donor plasmid-to-target plasmid transposition assay (**Fig. 1c**) (32,52). We compared WT PB to mutants with the three phosphorylated serine residues mutated to Ala (“AllStoA”) or individually to Pro (S17P, S35P, or S41P). We also tested the three Ser residues mutated to Glu (“AllStoE”) as possible mimics of phosphorylation. As shown in **Fig. 1d**, all mutations increased activity relative to WT, with the highest activity (4-fold increase) observed with the AllStoA mutant. Although all three individual Ser-to-Pro mutations contributed to the activity increase, the most important contributor appeared to be S35. AllStoA PB also exhibited increased activity as assessed by colony count assay using integration of a neomycin resistance transposon in human cells (**Fig. 1e**) (28).

### Predicted structure of full-length PB suggests that N-terminus phosphorylation inhibits DNA binding

To understand how the N-terminal region of PB might be regulating activity, we took advantage of the predictive power of AlphaFold-Multimer (37). Models were generated for the full-length PB dimeric transposase that were ranked by pLDDT score (local distance difference test) and, in all models, segments of the first 116 amino acids of PB folded with defined secondary structures that packed against the rest of the protein (**Fig. 2a,b**). Common to all models was an α-helix from Asp7-Leu15 pointing towards the transposon end binding site indicated in the cryo-EM structures. Also predicted was an α-helix (Ser76-Arg81) that, together with a short β-hairpin (β-strands Thr89-Gly92 and Lys95-Trp98), extended a β-stranded insertion domain in the transposase catalytic core thereby suggesting that they might participate in target DNA binding by increasing the target binding protein/DNA surface observed in the cryo-EM structure.

Of particular interest, several of the highest confidence models predicted that stretches of the first 116 residues are arranged *in trans*, i.e., residues from one PB monomer cross over and pack against the second monomer. In such an arrangement (**Fig. 2a**), all three N-terminal phosphorylated Ser residues were positioned such that they would conflict with DNA binding: S17 was positioned where bp 8 of the transposon end is located, and S35 and S41 were within the target binding site. This suggested the possibility that negatively charged phosphorylated N-terminal Ser residues may act as competitors to DNA binding, an effect that would be diminished by non-phosphorylated PB or by N-terminal truncation.

### Deletion of the N-terminal 74 residues of PB overcomes inhibition of transposition

To evaluate the potential of removing the observed inhibitory effect of phosphorylation, we generated two N-terminally truncated mutants of PB, Δ74PB and Δ104PB. To test the effect of N-terminal truncation, Δ74PB transposase was tested for transposon excision and integration in the LE-RE and LE-LE format (**Fig. 3a**), where excision was evaluated using a PCR-based excision assay and colony count assays served as proxy for integration of a neomycin-resistant (NeoR) transposon (**Fig. 3b**) (28). In the LE-LE format, WT PB exhibits little to no excision (**Fig. 3c, lane 5**), and hyPB manifests very low excision activity (**Fig. 3c, lane 14**); neither is capable of LE-LE integration (**Fig. 3d**). However, Δ74PB and Δ74hyPB are not only capable of LE-LE excision (**Fig. 3c, lanes 6 and 15**), they both exhibit higher integration activity than PB or hyPB with LE-RE transposons (**Fig. 3d**).

### Deletion of the N-terminal 104 AAs of PB results in a novel phenotype of excision active/integration inactive transposase on LE/LE transposon DNA

A particular advantage of using PB for genome engineering is its precise excision. Previous mutation of PB resulted in an excision active/integration inactive (exc+/int-) transposase that excises LE-RE transposons (19) but cannot reintegrate them. We considered the Alphafold2 structural prediction that AA 75-100 of PB may participate in target binding by extending the target binding protein/DNA surface observed in the cryo-EM structure (**Fig. 2a,b**). Therefore, we further truncated the N-terminal region beyond Δ74 to generate Δ104PB and evaluated its LE-LE excision and integration activity. In contrast to Δ74PB and satisfyingly consistent with AlphaFold predictions, Δ104PB was capable of excising but not integrating LE-LE transposon DNA into the human genome (**Fig. 4a,b**). Repair of the donor plasmid was unaffected as sequencing of the excision PCR product demonstrated precise TTAA reconstitution post-excision (**Fig. 4c**). Therefore, Δ104PB represents an exc+/int-transposase for LE-LE transposons.

### N-terminus is required for transposase dimerization prior to transposon end binding

Although we attempted to explore the role of the PB N-terminus using purified proteins, we were unable to recover any soluble N-terminally truncated proteins under conditions successfully used for WT PB in EXPI293F cells (26). To circumvent this, we introduced a TEV protease cleavage site between resides 74 and 75 which allowed us to express and purify a full-length protein and then remove the first 74 amino acids by cleavage after expression. Evaluation of oligomerization state by analytical ultracentrifugation revealed that, like WT, PB1-558 was a monodisperse dimer with a species at a sedimentation coefficient of 6.72 S corresponding to 125 kDa (93% of the absorbance signal; in blue, **Fig. 2c**). Conversely, PB74-539 generated by cleavage was a monomer with a dominant species at a sedimentation coefficient of 3.58 S corresponding to 53.4 kDa (88% of the absorbance signal; in red, **Fig. 2c**). This result supported the *in trans* models for PB dimerization and is consistent with observations that full length PB is a dimer even prior to transposon end binding and synapse (14,26). We suggest that the *trans* binding segment of the N-terminus is displaced upon transposon end binding and after synapse formation the extensive network of protein-DNA interactions seen in the cryo-EM structures now hold the dimer together; this would explain why the N-terminal region was disordered in the cryo-EM maps.

### Addition of a second C-terminal domain to PB overcomes the lack of activity on symmetric transposon ends

Our recent cryo-EM structures of the dimeric PB transposase bound to two oligonucleotides representing two transposon LEs showed that the multidomain enzyme recognizes motifs within its ends in a modular fashion (26). The tip of each transposon end is bound by the central catalytic domain (residues 264-456), and a DNA motif just inside the transposon ends (green, **Fig. 1a)** is recognized by a bipartite DNA binding domain (residues 117-263 and 457-535). However, the strongest binding affinity is conferred by the cysteine-rich C-terminal domain (“CD”; residues 553-594) that dimerizes as it binds to a 19-bp palindrome (purple). Although both transposon ends have these palindromes, given that the dimeric PB transposase can supply only two CDs, the surprising result was an asymmetric structure (**Fig. 5a**) with the palindrome of one LE bound by two CDs while the palindrome of the other LE was bound by nothing and was disordered in the electrostatic potential density map. As the palindromes are asymmetrically arranged on the LE and the RE terminal inverted repeats (TIRs) (**Fig. 1a**), the most straightforward model of a transpososome assembled on the active LE-RE ends is a tetramer (**Fig. 5b**) that could provide the two CDs to each of the two palindromes (26). As PB does not excise or integrate a transposon with symmetrical LE-LE ends in cells (26), this tetramer model is very likely describing the active transpososome architecture.

The tetrameric LE-RE transpososome implies that appending a second copy of residues 543-594 immediately following F594 (**Fig. 5b**), a design that would supply two CDs to the 19-bp palindrome on LE, should permit transposition using symmetrical LE-LE transposon ends in cells. The architecture of PB transpososomes indicated that the addition of a second CD (2CD) could be accommodated since a long, partially disordered linker connects residues 535 and 553 of the CD, and there are no protein-protein interactions between either of the CDs and the rest of the transposase (**Fig. 5a**). When we tested this new transposase in human cells, the effect on excision and integration by PB-2CD and hyPB-2CD was very similar to that observed for Δ74: both were active on LE-LE transposons (**Fig. 3c, lanes 7 and 16; Fig. 3d)**, and rescued inactive PB and hyPB excision and integration of LE-LE transposons (**Fig. 3c, lanes 7 vs. 5 and 16 vs. 14; Fig. 3d**).

**Fig. 3.**
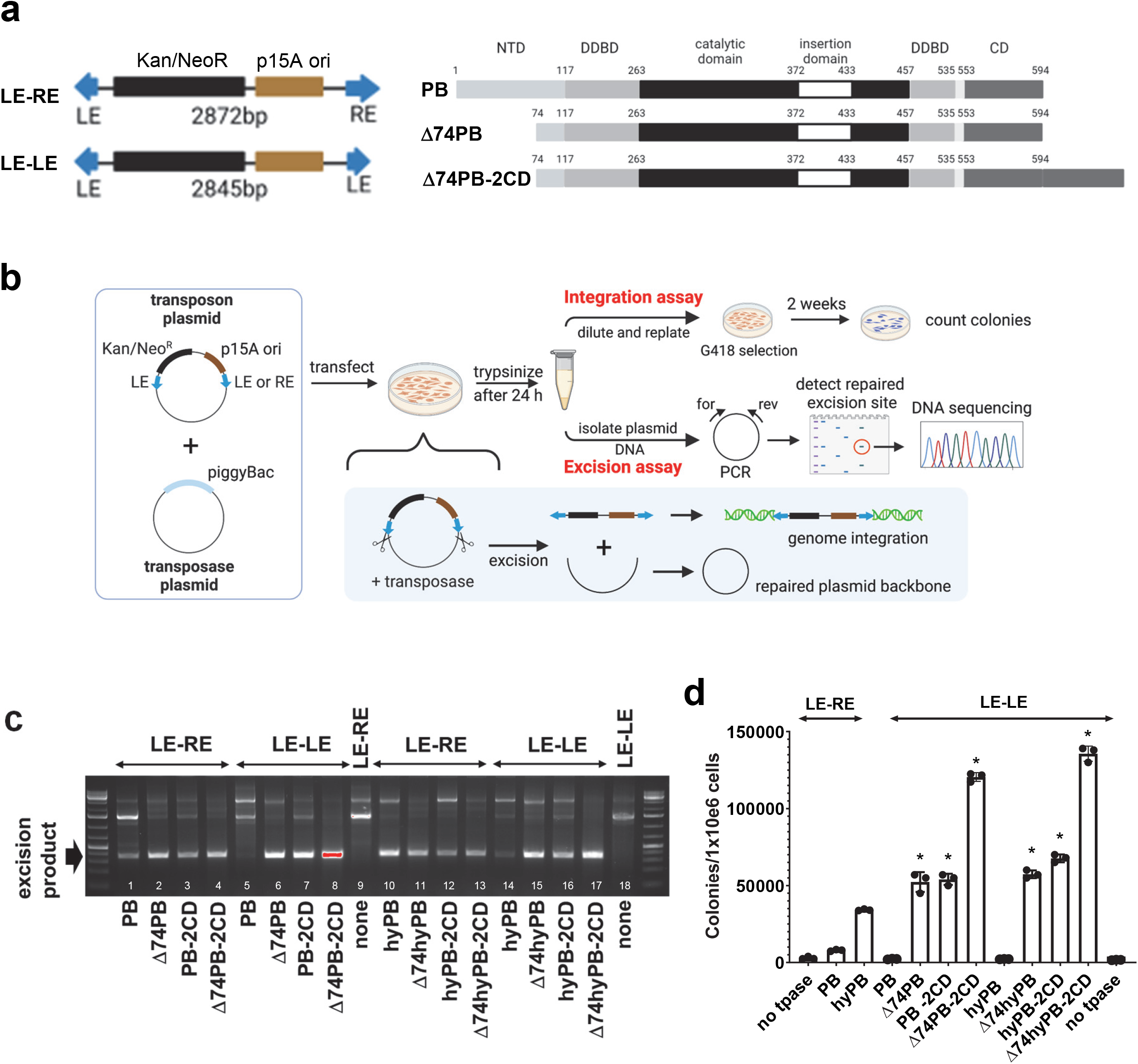
Redesigned PB overcomes inhibition in human cells. **a**. Left, schematic of LE-RE vs LE-LE transposons containing a kanamycin/neomycin resistance cassette (Kan/NeoR) and a p15A origin of replication (p15A ori). Right, schematics of WT PB compared to Δ74PB and Δ74PB-2CD. The catalytic domain contains the conserved DDD motif. NTD N-terminal domain. Dimerization and DNA-binding domain (DDBD). CD, C-terminal cysteine-rich domain. **b**. Schematic of transposition in human cells evaluated via transposon excision and integration (colony count) assays. **c**. Excision assay analysis of Δ74PB and Δ74PB-2CD with LE-LE compared to WT PB and hyPB with LE-RE. Shown is representative of three independent experiments. **d**. Colony count (integration) assay analysis of Δ74PB and Δ74PB-2CD with LE-LE compared to WT PB and hyPB with LE-RE. N=3±SEM; *, p<0.05 compared to PB or hyPB respectively with LE-RE using one way ANOVA and Turkey multiple comparisons test.

**Fig. 4.**
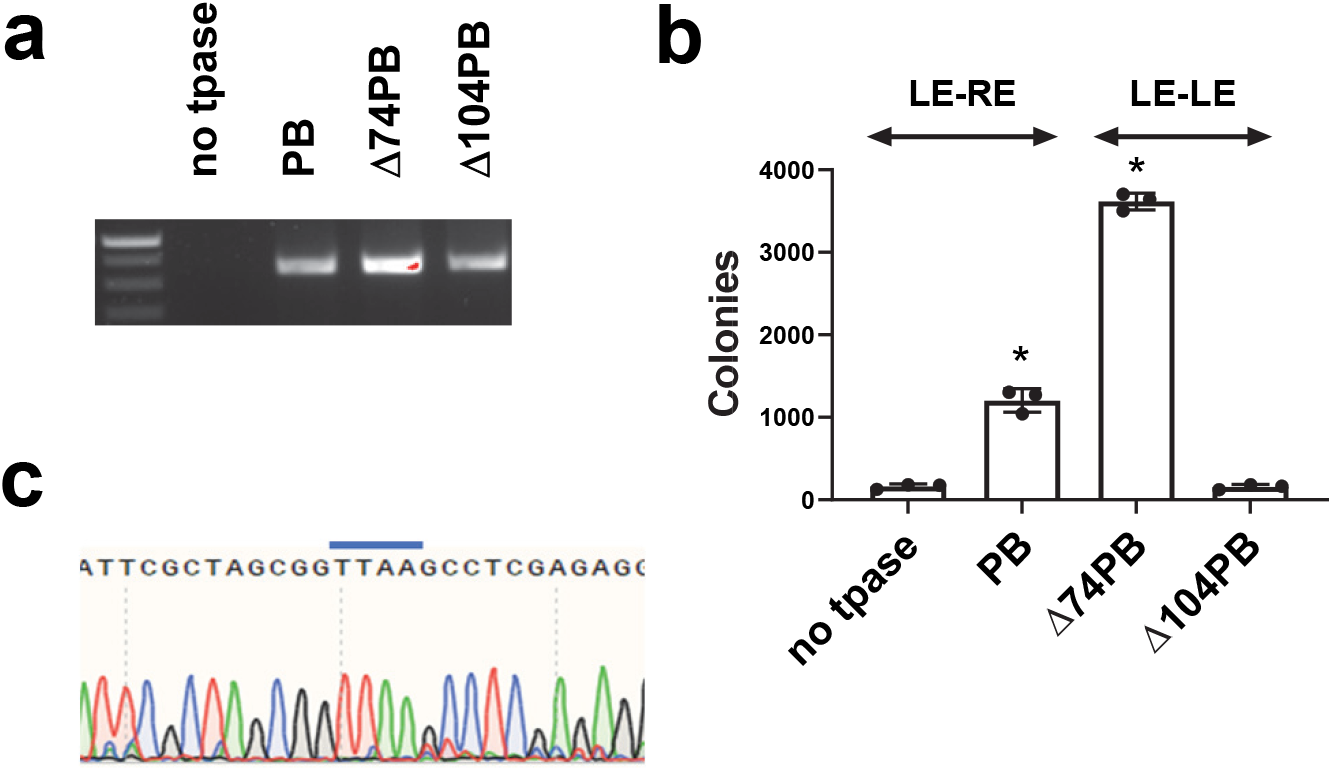
Δ104PB is an excision active/integration inactive transposase on symmetric LE-LE TIRs. **a**. Excision assay analysis of Δ104PB compared to WT and Δ74PB in human cells. Shown is representative of 3 independent experiments. **b**. Colony count (integration) analysis of Δ104PB compared to WT and Δ74PB in human cells. N=3±SEM; *, p<0.05 using one way ANOVA and Dunnett’s multiple comparisons test compared to no transposase control. **c**. Sanger sequencing of excision PCR products demonstrating TTAA reconstitution with Δ104PB.

**Fig. 5.**
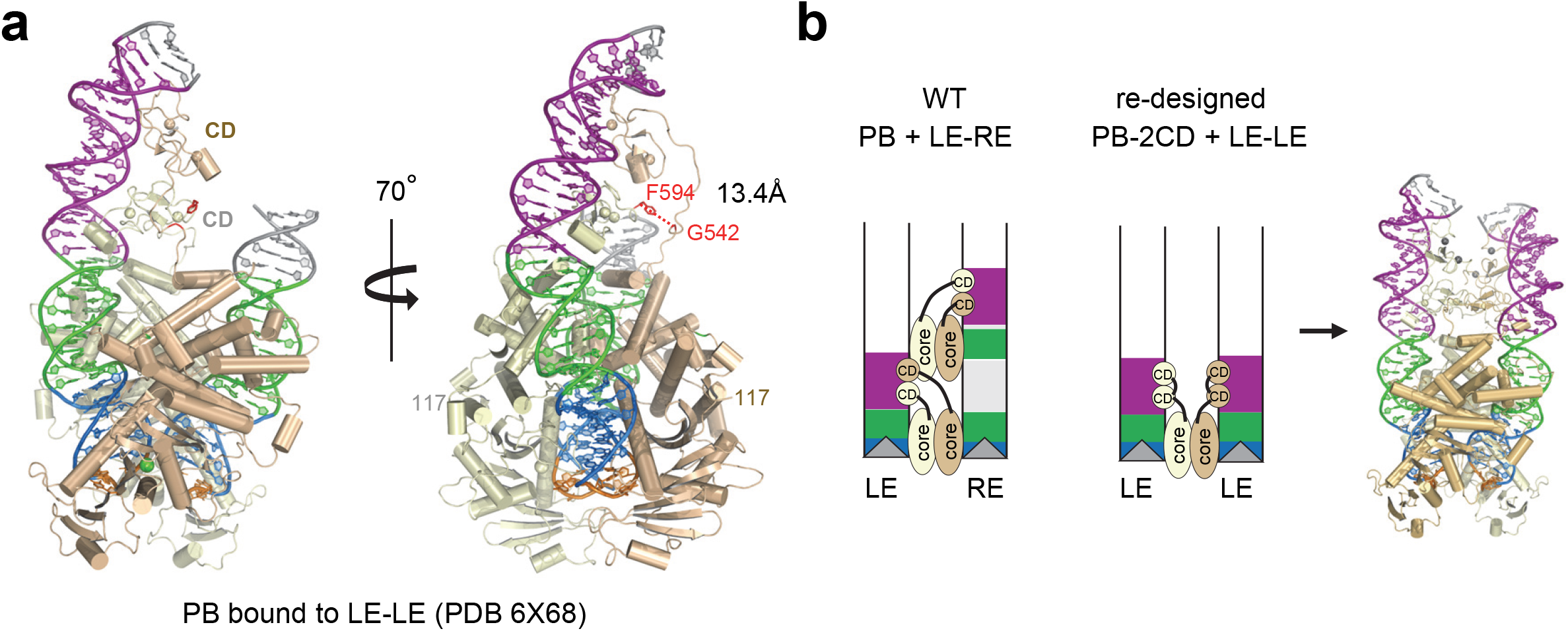
Structure based re-design of PB for symmetric transposon ends. **a**. Structure of PB bound to LE-LE transposon. **b**. Model of asymmetric PB tetramer bound to LE-RE transposon. Re-design permits symmetric PB dimer to bind LE-LE transposon via appending a 2CD to the end of the PB transposase.

### Δ74PB-2CD overcomes inhibition over each modification alone

When we combined the two transposase modifications, both Δ74PB-2CD and Δ74hyPB-2CD exhibited high excision activity (**Fig. 3c, lanes 8 and 17**) and additive integration activity when compared to Δ74- or -2CD transposases alone with LE-LE transposons (**Fig. 3d**). We also evaluated excision and integration activities of the two most active transposases, Δ74PB-2CD and Δ74hyPB-2CD, over a range of transposase doses (5ng to 2.5µg) in combination with a fixed amount of transposon DNA (1.5µg) (**Fig. 6a, b**). At 500ng of transposase DNA, we observed a dramatic 20-fold increase in integration efficiency by Δ74PB-2CD with LE-LE relative to PB with LE-RE (**Fig. 6b**). Even when compared with hyPB with LE-RE, at that same transposase dosage, we observed 5-fold increased integration efficiency for Δ74hyPB-2CD with LE-LE (**Fig. 6b**).

**Fig. 6.**
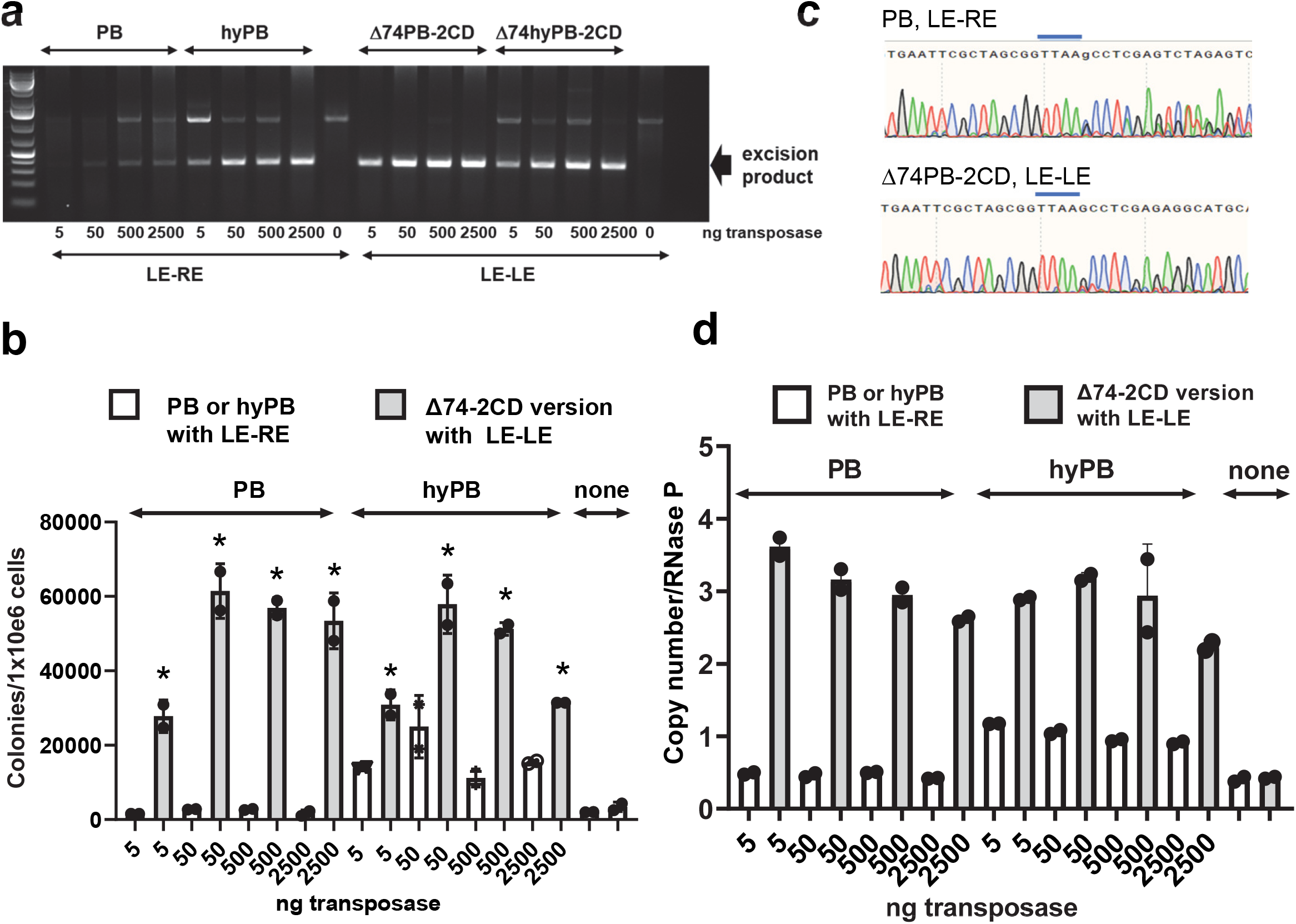
Redesigned PB overcomes inhibition over a range of transposase doses in human cells. **a**. Excision assay analysis of Δ74PB and Δ74PB-2CD with LE-LE compared to WT PB and hyPB with LE-RE over a range of transposase dosages while keeping transposon DNA constant at 1.5µg. Shown is representative of 3 independent experiments. **b**. Colony count (integration) analysis corresponding to the excision analysis in a. N=3±SEM*, p<0.05 compared to PB or hyPB respectively with LE-RE using one way ANOVA and Šídák’s multiple comparisons test. **c**. Sanger sequencing of excision PCR products comparing WT PB with LE-RE to Δ74PB-2CD with LE-LE demonstrating TTAA site reconstitution. **d**. ddPCR copy number analysis of the number of integrated transposons in human cells normalized for the RNaseP gene. N=2±SEM.

We confirmed that Δ74PB-2CD with LE-LE transposon ends reconstituted the TTAA target site after excision by sequencing the excision PCR product, thus validating the key molecular signature of PB-mediated excision (**Fig. 6c**). We determined the copy number of integrated transposons in human cells when using 1µg of transposase with 1.5 µg of transposon DNA. As expected and consistent with excision and integration analysis, there were more integration events for Δ74PB-2CD or Δ74hyPB-2CD with LE-LE transposons than with PB or hyPB with LE-RE transposons (**Fig. 6d**). Δ74PB-2CD or Δ74hyPB-2CD also outperformed their counterparts in integrating a range of small to very large transposons (3.4kb, 7.8kb, and 15.1kb) in human cells (**Fig. 7**), indicating that these modifications overcome the negative regulation mediated by N-terminal phosphorylation and asymmetric transposon ends.

**Fig. 7.**
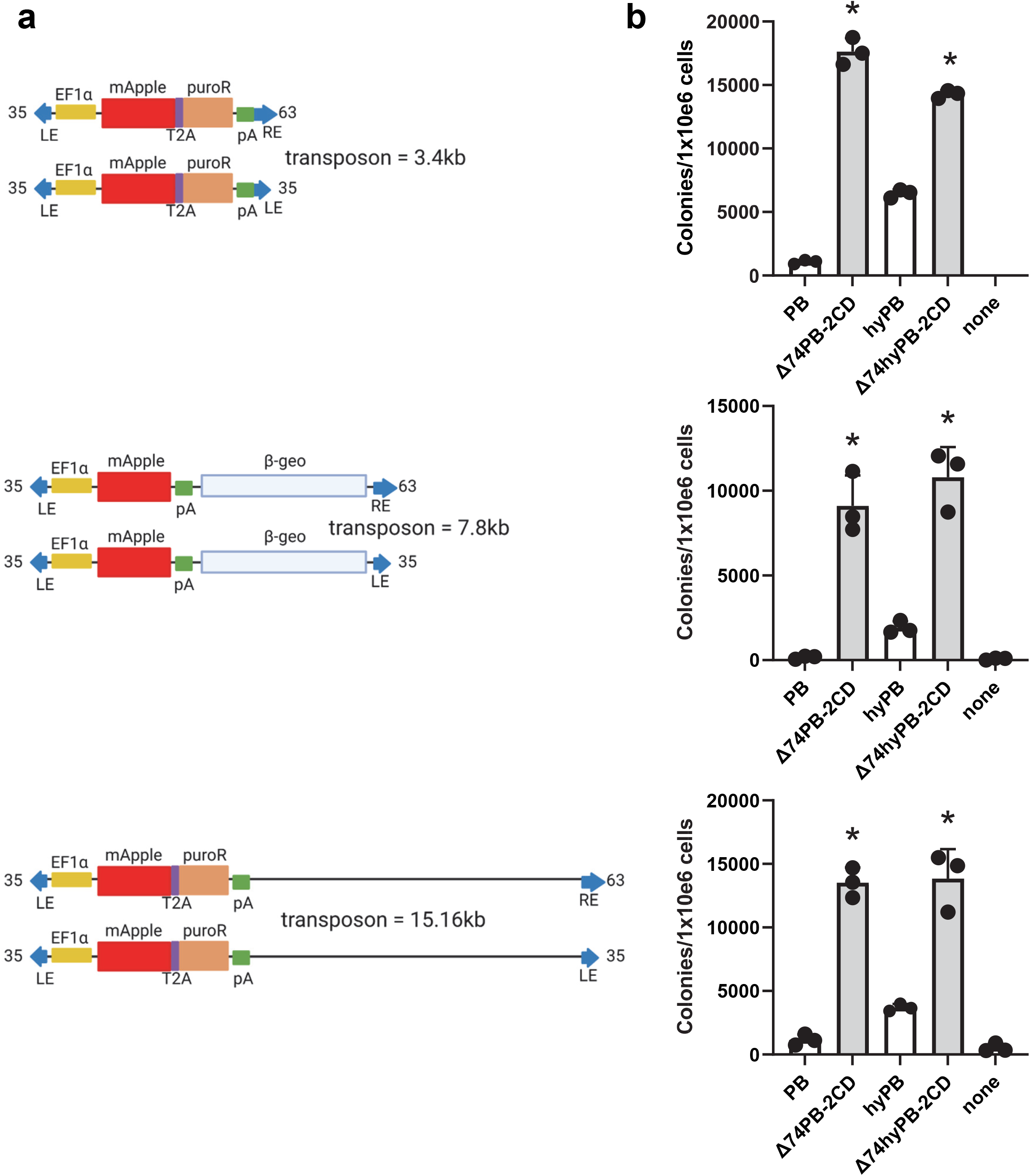
Redesigned PB overcomes inhibition over a range of transposon sizes in human cells. **a**. Schematics of transposon vectors of varying sizes ranging from 3.4 to 15.1kb. **b**. Colony count (integration) analysis corresponding to transposon sizes in **a**. N=3±SEM; *, p<0.05 compared to PB or hyPB respectively with LE-RE using one way ANOVA and Turkey multiple comparisons test using the transposons in **a**.

### Δ74PB-2CD demonstrates an unaltered integration profile in human cells

Some previous modifications to PB have resulted in loss of integrity of precise excision and target site duplication (53). To evaluate the integration site profile of our modified PB transpososomes, we used self-reporting transposon technology developed by Moudgil et al. to evaluate for potential alterations in integration site profile in HCT116 cells (30). We observed precise and consistent excision and target site duplication for all transpososomes tested (**Fig. 8a, S2a and b**). Our genome-wide analyses of integration sites revealed a broad and consistent distribution across all chromosomes for WT and redesigned transpososomes (**Fig. 8b, S2c**). Further annotation by genomic features showed comparable preferences for different genomic regions for all PB transpososomes (**Fig. 8c, S2d**). To characterize individual insertion sites in more detail, we split the genome into one mega-base regions and plotted the normalized abundance of insertion sites for WT and PB mutants (**Fig. S3**). A candidate genomic region revealed similar insertion patterns for PB WT and mutants (**Fig. 8d**). Overall, our genome-wide analysis of insertion sites in human cells revealed that all redesigned PBs maintained precise excision and target site duplications.

**Fig. 8.**
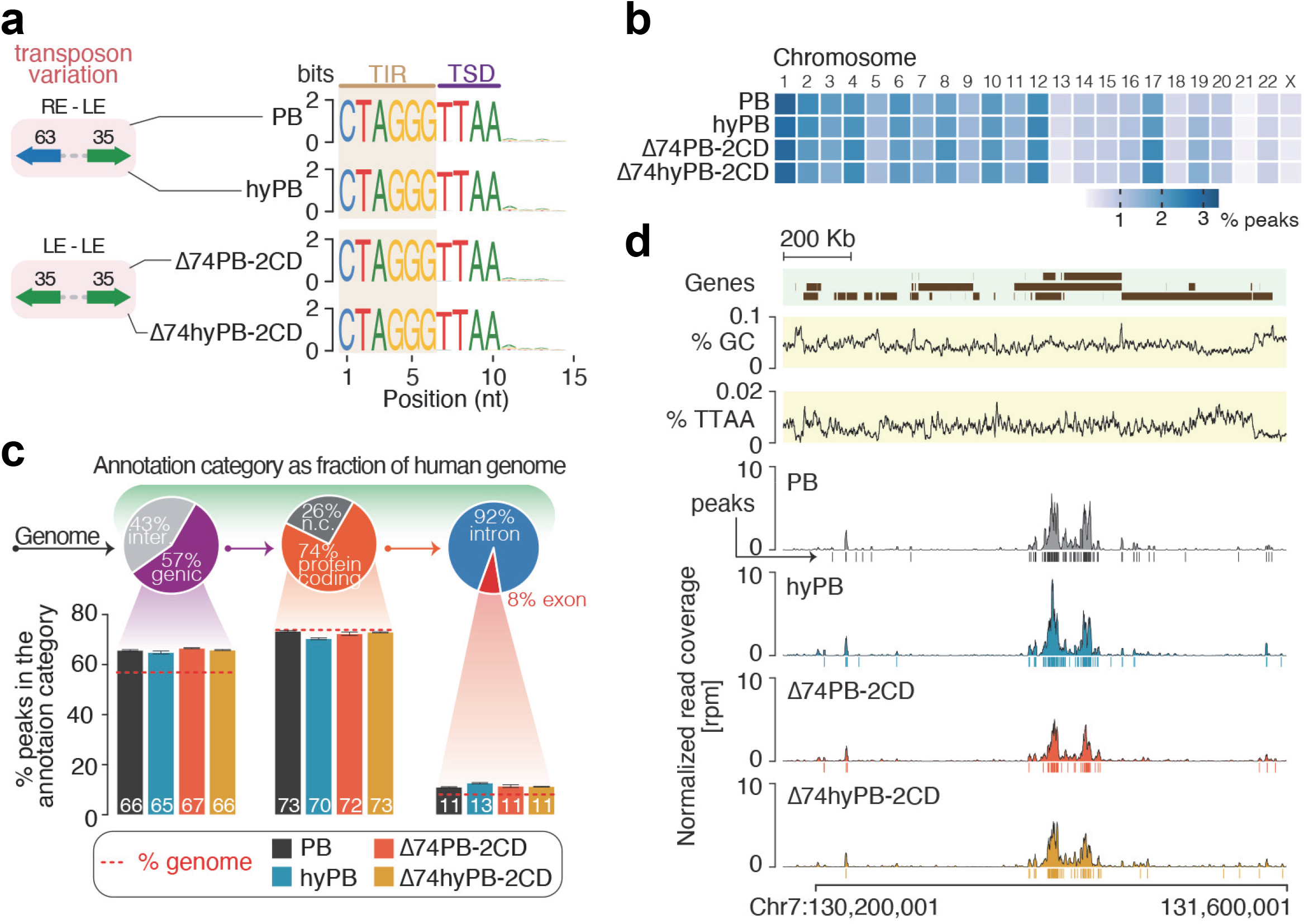
Genome-wide characterization of insertions sites by re-designed transposomes. **a**. The sequence logo of the 5’insertion site (first 15 nucleotides) including the transposon inverted repeat (TIR) and the target site showed consistent excision and target site duplication (TSD) for wild-type PB, hyperactive mutants, and different TIRs. Three biological replicates were analyzed, and a single representative replicate is shown. **b**. The overall genome-wide distribution of insertion-peaks remained unchanged between wild-type and mutant transposomes across different chromosomes [mean value, N=3]. **c**. Annotation of insertion-peaks by genomic features showed comparable preferences for different genomic regions [mean±SEM, N=3]. Genic versus intergenic regions, protein coding genes, and protein coding exons are depicted. The genomic contribution of these regions is indicated as pie charts for comparison. **d**. Insertions-peaks for wild type and mutant PB transposomes are shown for a representative genomic region with high-density insertions on Chromosome 7. Insertion densities are shown as normalized read coverage (rpm), and peaks are indicated.

## Discussion

The growing list of high-resolution three-dimensional structures of DNA transposases in complex with the specific DNA upon which they act has provided unprecedented insight into their mechanisms of action (54-57). For PB, for example, the observation that recognition of the transposon hairpin intermediate structurally mimics target site binding revealed an unanticipated link between the mechanisms of excision and integration (26).

Here, we have attained more mechanistic insight into PB transposition in mammalian cells using the AlphaFold system (37,58) in combination with cellular and biochemical assays. AlphaFold suggested several possible functional roles for the N-terminal 116 amino acids of PB which were disordered in the cryo-EM structures. Once we identified CKII phosphorylation sites within this region of PB and verified that phosphorylation in mammalian cells inhibits transposition, these data taken together suggested that CKII phosphorylation may interfere with the binding of transposon ends and target DNA, mechanistically explaining the inhibitory effect. Transposase phosphorylation has been reported previously; for example, Protein Kinase A dependent inhibition of Mos1 has been observed (59), although only a small fraction of the Mos1 transposase appears phosphorylated in S2 insect cells. PB phosphorylation at multiple sites demonstrates novel involvement of CKII in downregulating a mobile genetic element. CKII is a constitutive kinase primarily, but not exclusively located in the nucleus (60), and phosphorylation of a transposase by a constitutive kinase to inhibit DNA binding seems a clever way to tame a mobile element. Therefore, transposon downregulation can now be added to the extraordinary pleiotropy of CKII. Perhaps phosphorylation could be used to regulate PB activity and timing of transposition in mammalian cells.

The AlphaFold predictions also provided a framework for understanding our observation that Δ104PB is exc+/int-transposase on LE-LE transposon DNA. This deletion mutant of PB may be particularly valuable if Δ74PB-2CD or Δ74hyPB-2CD are used for high efficiency transposon integration, as those LE-LE transposons can still be excised by Δ104PB if needed in downstream applications. This therefore complements previous mutational result for PB that resulted in an exc+/int-transposase that excises LE-RE transposons (19) but cannot reintegrate them. In fact, it appears that the combination of AlphaFold with experimental structures is particularly powerful as it can provide hypotheses as to the function of parts of molecular assemblies that are invisible in experimental structures due to disorder.

An example of the power of this synergistic relationship between experiment and prediction was a question left unanswered by our cryo-EM structures. Biophysical data indicated that PB is a dimer when expressed in mammalian cells (14), even before the formation of synaptic complex, yet the almost entirely polar protein-protein interface between the two monomers is too small at 644 A^2^ and did not explain dimerization prior to DNA binding. AlphaFold prediction suggested that the phosphorylated N-terminus might provide additional monomer-monomer interactions prior to DNA binding thus promoting dimerization, and we have confirmed this using purified proteins and sedimentation analysis. We now suspect that, upon synaptic complex formation, the bound transposon ends may displace the N-terminus while maintaining transposase dimerization.

We have shown the native PB transpososome can be rationally simplified to a symmetric dimer via the addition of a second CD to the transposase resulting in high activity on symmetric LE-LE transposon ends upon which WT PB is inactive, an effect further increased by the Δ74 truncation. This redesign of the PB transpososome to include Δ74PB-2CD in combination with symmetric LE-LE ends has produced a novel highly active transpososome with apparently unaltered excision and integration fidelity. Our redesigned PB transpososomes exhibit unaltered integration site profile, no loss integrity of transposon ends or target site duplication, and offer the ability to excise and not re-integrate transposon DNA if needed. These simplified PB transpososomes provide novel scaffolds that may allow for further engineering to achieve other outcomes such as high efficiency targeted transposon integration in mammalian cells which thus far has remained elusive. In particular, the ability to reduce the number of appended targeting domains from four in a homotetrameric PB transpososome to only two in a dimeric Δ74PB-2CD transpososome may be a particularly attractive benefit.

One aspect that unites otherwise surprisingly diverse transposome architectures is their modularity. It appears that the RNAseH-like core can accommodate a variety of different insertions of other domains and variable numbers of different DNA binding domains (often Zn finger variants) fused to it. These architectures, if characterized precisely by three-dimensional structures, could provide mechanistic insight for other transpososomes. Modularity offers re-engineering possibilities. Furthermore, computational tools such as Rosetta are developing rapidly that allow the design of new protein/protein interfaces to create stable assemblies with novel functions (61). One envisions a future in which the combination of experimental and computational tools will be used to generate novel DNA transpososomes to add to the genomic tool kit available for genomic and clinical applications.

## Supporting information

Supplemental computation methods

Supplemental mass spec table

## Data availability

All primary data is available from the authors upon request. All sequencing data generated in this study has been deposited at GEO and is publicly available under accession GSE201914.

## Code availability

All relevant code is provided in the supplemental methods and available on github website (https://github.com/HaaseLab/piggyBac_mutants). For additional information contact Astrid Haase.

## Acknowledgments

M.H.W. was supported by National Institutes of Health R01 DK093660, R01 EB033676, and P30 DK114809. F.D. (ZIA DK036153-16) and A.H. (1ZIADK075111-07) were supported by Intramural Program of the National Institute of Diabetes and Digestive and Kidney Diseases of National Institutes of Health.

