## Supplemental computation methods for "Transposase N-terminal phosphorylation and asymmetric transposon ends inhibit *piggyBac* transposition in mammalian cells"

Computational methods by Pavol Genzor

***Wentian Luo, Alison B. Hickman, Pavol Genzor,*** *Rodolfo Ghirlando,* ***Christopher M. Furman, Anna Menshikh, Astrid Haase, Fred Dyda, and Matthew H. Wilson***

This document contains sample of computational methods associated with the manuscript. Please contact Dr. Astrid Haase with any questions.

Related code, sample data and functions are also available at github https://github.com/HaaseLab/piggyBac_mutants page.

**Table of Content**


- Raw data processing
  - Read trimming and genomic alignments parameters
- ANALYSIS
  - Integration and target site duplication analysis
  - Loading paired-end (PE) .bam into R
  - Identifying integration regions - “peaks”
- PLOTS
  - Chromosome peak distribution
  - Genome annotation
  - Tiled integration map
  - Genome coverage
- Functions
  - filterBamPE()
  - findPeaksInPERegions()

**Read trimming and genomic alignments parameters.**

```
## FIRST trimming to remove sequencing adapters

cutadapt  -n 4 -j 6 -m 26
          --pair-filter=any --discard-untrimmed
          -g AGATGTGTATAAGAGACAG -a AAAGATAGTCTGCGTAAA -G TTTACGCAGACTATCTTT -A CTGTCTCTTATACACATCT
          -o R1_output_path -p R2_output_path
          R1_input.fastq.gz  R2_input.fastq.gz

## SECOND trimming to remove the TIR sequences

cutadapt  -n 1 -j 6 -m 20
          --pair-filter=both --discard-untrimmed
          -a CCCTAG -G CTAGGG
          -o R1_output_path -p R2_output_path
          R1_input.fastq.gz R2_input.fastq.gz

## GENOMIC ALIGNMENT

hisat2  -p 6 --dta --rna-strandness RF
        -x /data/genzorp/genomes/GenCode/HiSat2Index_h38.p5/GRCh38.p5.v24
        -1 R1_second_trim.fastq.gz -2 R2_second_trim.fastq.gz | samtools sort -@ 6 -m 2G - -o
        sample_name.bam
```

**Integration and target site duplication analysis**

Reads containing the TIR sequence are loaded into R and the the nucleotide frequencies for the first fifteen nucleotides of the reads containing TIR are calculated.

```
## libraries
library(data.table);library(parallel);library(GenomicRanges);library(ShortRead)

## session dir
SESSION.DIR="/data/Rsessions/"

## TRIM ONE
t1_fastq_dir <- "/data/Cutadapt/Trim_one/"
t1_fastq_file_names <- grep("R2",list.files(t1_fastq_dir),value=TRUE)
an_extention <- "_R2_trimmed.fastq.gz"
t1_fastq_sample_names <- gsub(an_extention,"",t1_fastq_file_names)
t1_fastq_file_paths=paste0(t1_fastq_dir,t1_fastq_file_names)
names(t1_fastq_file_paths) <- t1_fastq_sample_names
t1_fastq_file_paths

## load fastq files
FQL <- lapply(names(t1_fastq_file_paths),function(s){
  message(paste0("loading: ",s))
```

```
  fq <- readFastq(dirPath=t1_fastq_file_paths[[s]])
  return(fq)})
names(FQL) <- names(t1_fastq_file_paths)

## Get specific names
pBac_names <- names(FQL)

## All data logo
bac.all.NFL <- lapply(pBac_names, function(s){
  message(paste0("sample: ",s))
  aSeq <- sread(FQL[[s]])
  aSeq.sub <- subseq(x = aSeq, start = 1, width = min(width(aSeq)))
  abc.mat <- alphabetByCycle(stringSet = aSeq.sub, alphabet = c("T","C","A","G"))
  return(abc.mat)}); names(bac.all.NFL) <- pBac_names

##
## SELECT SUBSETS with TIR only
##

## search for sequence
bac_tir <- "^CTAGGG"

## ONLY TIR logo
bac.tir.NFL <- lapply(pBac_names,function(s){
  message(paste0("sample: ",s))
  aSeq <- sread(FQL[[s]])
  aSeq.tir.short <- aSeq[grep(bac_tir,aSeq)]
  aSeq.tir.short.sub <- subseq(x = aSeq.tir.short, start = 1,
                               width = min(width(aSeq.tir.short)))
  abc.mat <- alphabetByCycle(stringSet = aSeq.tir.short.sub,
                             alphabet = c("T","C","A","G"))
  return(abc.mat)}); names(bac.tir.NFL) <- pBac_names
```

Frequencies were plotted to show any changes in the integration preferences using sequence logo.

```
## LOAD DATA
load("/Users/genzorp/Documents/GITHUB/LIVE/piggyBac_mutants/data/PB_freqTables.RData")
FAM="Helvetica"; XYT=8; TCOL="black"

## View data
## bac.all.NFL; bac.tir.NFL

## Settings
suppressPackageStartupMessages(library(ggseqlogo))
NUC.COLORS <- make_col_scheme(chars = c("T","C","G","A"),
                              cols = c("firebrick1","dodgerblue1","goldenrod1","forestgreen"))

## Sample order
bac_order <- c("HCT116_re67_le40_PB_1","HCT116_re67_le40_PB_2","HCT116_re67_le40_PB_3",
               "HCT116_re67_le40_hyPB_1","HCT116_re67_le40_hyPB_2","HCT116_re67_le40_hyPB_3",
               "HCT116_le40_le40_delta74PB2CD_1", "HCT116_le40_le40_delta74PB2CD_2",
```

```
                "HCT116_le40_le40_delta74PB2CD_3", "HCT116_le40_le40_delta74hyPB2CD_1",
                "HCT116_le40_le40_delta74hyPB2CD_2", "HCT116_le40_le40_delta74hyPB2CD_3")

##
## Tabulate number or reads and calculate percentage with TIR
##

bac.rcdt <- data.table("samples"= names(lapply(bac.all.NFL,function(s){sum(s[,1])})),
                       "input_reads"= unlist(lapply(bac.all.NFL,function(s){sum(s[,1])})),
                       "tir_reads" = unlist(lapply(bac.tir.NFL,function(s){sum(s[,1])})))
bac.rcdt[,"percent_with_tir":=round((tir_reads/input_reads)*100,3)]
bac.rcdt
```

```
##                               samples input_reads tir_reads percent_with_tir
##                                <char>       <int>     <int>            <num>
##  1: HCT116_le40_le40_delta74hyPB2CD_1      391684    381248           97.336
##  2: HCT116_le40_le40_delta74hyPB2CD_2      382976    372018           97.139
##  3: HCT116_le40_le40_delta74hyPB2CD_3      421934    410068           97.188
##  4:   HCT116_le40_le40_delta74PB2CD_1      380435    371180           97.567
##  5:   HCT116_le40_le40_delta74PB2CD_2      390375    380870           97.565
##  6:   HCT116_le40_le40_delta74PB2CD_3      410274    400915           97.719
##  7:           HCT116_re67_le40_hyPB_1      437558    425128           97.159
##  8:           HCT116_re67_le40_hyPB_2      419371    406653           96.967
##  9:           HCT116_re67_le40_hyPB_3      443495    429986           96.954
## 10:             HCT116_re67_le40_PB_1      344284    337004           97.885
## 11:             HCT116_re67_le40_PB_2      349753    340907           97.471
## 12:             HCT116_re67_le40_PB_3      388462    378610           97.464
```

```
##
## Plot Sequence LOGOS with TIR
##

bac.tir.logo <- suppressWarnings(suppressMessages(
  lapply(bac_order,function(s){
    aMat <- bac.tir.NFL[[s]]
    aTitle <- paste0(s,"\n","TIR reads = ",
                     bac.rcdt[samples %in% s][["tir_reads"]],"\n", "% with TIR = ",
                     bac.rcdt[samples %in% s][["percent_with_tir"]])
    ggL <- ggseqlogo(data = aMat, font = "helvetica_regular", col_scheme = NUC.COLORS)
    ggLplus <- ggL + theme_void() + ggtitle(aTitle) +
      scale_x_continuous(breaks = c(1,seq(0,100,5))) +
      xlab("Position (nucleotides)") +
      theme(aspect.ratio = 0.2, axis.text = element_text(family = FAM, color = TCOL),
            axis.title = element_text(family = FAM, colour = TCOL),
            axis.title.y = element_text(angle = 90),
            axis.ticks = element_line(colour = TCOL), axis.ticks.length = unit(2,"mm"))
    return(ggLplus)})))
names(bac.tir.logo) <- bac_order

## Print a single plot
bac.tir.logo$HCT116_re67_le40_PB_1
```

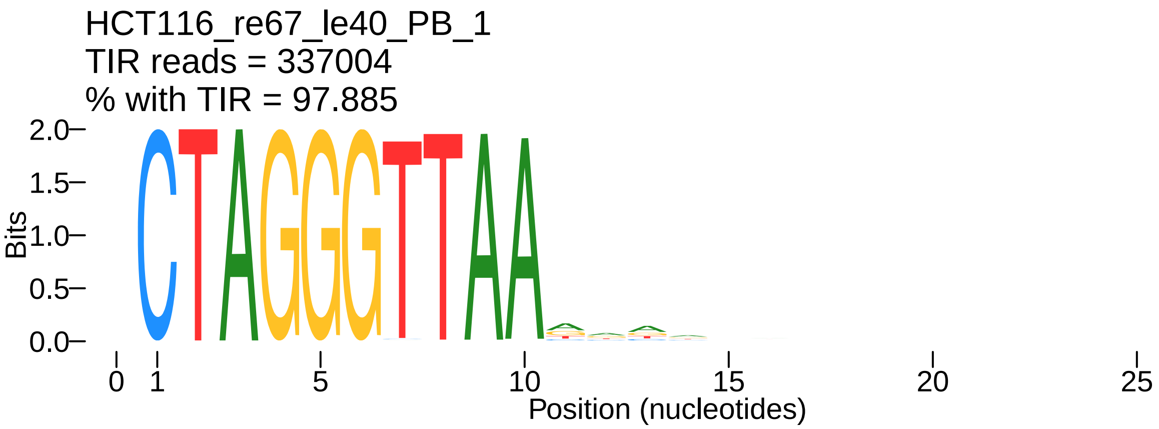


**Loading paired-end (PE) .bam into R environment**

First, paired-end *.bam* files are loaded into R and paired end reads are converted into unique genome mapping fragments using the custom function *filterBamPE*. The function is at the end of document and available online on

```
## NOTE: THIS CAN TAKE A VERY LONG TIME AND MEMORY
## NOTE: Make sure to use small files or have enough resources

## function
library(parallel)
source("/Users/genzorp/Documents/GITHUB/LIVE/piggyBac_mutants/r/filterBamPE.R")

## directories
SESSION.DIR="/Users/genzorp/Documents/GITHUB/LIVE/piggyBac_mutants/sessions/"

## files
BAM.DIR.TRIM.TWO="/Users/genzorp/Documents/GITHUB/Hisat/Trim_two/"
BAM.FILE.NAMES <- grep("bam$",list.files(BAM.DIR.TRIM.TWO), value = TRUE)
BAM.SAMPLE.NAMES <- gsub(".bam","",BAM.FILE.NAMES)
BAM.FILE.PATHS <- paste0(BAM.DIR.TRIM.TWO,BAM.FILE.NAMES)
names(BAM.FILE.PATHS) <- BAM.SAMPLE.NAMES
BAM.FILE.PATHS

## settings
STRAND.MODE=2
BS.SPECIES="Hsapiens"
MANUAL.WIDTH.FILTER=1000

## Load reads into R
BPEL <- lapply(names(BAM.FILE.PATHS), function(s){
  message(paste0("processing: ",s))
  filterBamPE(BAM.FILE = BAM.FILE.PATHS[[s]],
              BAM.NAME = s,
              STRAND.MODE = STRAND.MODE,
              BS.SPECIES = BS.SPECIES)})
names(BPEL) <- names(BAM.FILE.PATHS)
```

```r
## Create sample table
dtsi <- data.table("original"=names(BPEL),
                   "te1"=unlist(tstrsplit(names(BPEL), split = "_", keep = 2)),
                   "te2"=unlist(tstrsplit(names(BPEL), split = "_", keep = 3)),
                   "genotype"=unlist(tstrsplit(names(BPEL), split = "_", keep = 4)),
                   "rep"=unlist(tstrsplit(names(BPEL), split = "_", keep = 5))); dtsi

bac_order <- c("HCT116_re67_le40_PB_1","HCT116_re67_le40_PB_2","HCT116_re67_le40_PB_3",
               "HCT116_re67_le40_hyPB_1","HCT116_re67_le40_hyPB_2","HCT116_re67_le40_hyPB_3",
               "HCT116_le40_le40_delta74PB2CD_1", "HCT116_le40_le40_delta74PB2CD_2",
               "HCT116_le40_le40_delta74PB2CD_3","HCT116_le40_le40_delta74hyPB2CD_1",
               "HCT116_le40_le40_delta74hyPB2CD_2", "HCT116_le40_le40_delta74hyPB2CD_3")

all_order <- bac_order
all_order

## clean session
keep_objects <- c("BPEL","dtsi","all_order","bac_order")
remove_objects <- setdiff(ls(),keep_objects)
rm(list = remove_objects)

## save
#save.image(file = "/Users/sessions/loadedBams.RData")
```

#### Identifying integration regions - “peaks”

Next, loaded fragments are used to find “peaks” - regions containing minimum number of unique sequences (min = 5) to be considered “good” integration sites.

```r
## NOTE: load data if not present
## load(file = "/Users/sessions/loadedBams.RData")

## NOTE: calculate coverage and find peaks
## PEAKS: to find peaks
##   a) calculate coverage
##   b) slice the coverage keeping regions > 5 reads
##   c) convert and reduce the sliced RleViews into GR
##   d) add area under the curve for each peak
##   e) count sequences within the region
##   f) add library normalized area under the curve (nauc)


## FIND PEAKS
## - NOTE: only sequences
source("/Users/genzorp/Documents/GITHUB/LIVE/piggyBac_mutants/r/findPeaksInPERegions.R")
peak.SL <- lapply(names(BPEL), function(s){
  message(paste0("sample: ",s))
  peakGR <- findPeaksInPERegions(IN.GR = BPEL[[s]],
                                 MIN.READS = 5,
                                 GR.NAME = s)
```

```r
  return(peakGR)})
names(peak.SL) <- names(BPEL)

## Export peak list as bed (for GEO)

#names(peak.SL)
#lapply(names(peak.SL), function(s){
#  agr <- peak.SL[[s]]
#  adt <- as.data.table(agr)
#  adt[,"peakN" := paste0("peak",1:nrow(adt))]
#  abed <- adt[,.SD,.SDcols = c("seqnames","start","end","peakN","peak_auc",
#                                "strand","peak_height","peak_width","total_regions")]
#  write.table(file = paste0(TAB.DIR,s,"_peaks.bed"), x = abed, quote = FALSE,
#              sep = "\t", row.names = FALSE, col.names = FALSE)})

## Add number of peaks to sample info table (dtsi)
dtsi[["peak_count"]] <- unlist(lapply(dtsi[["original"]], function(s){length(peak.SL[[s]])}))


## Record fraction of the usable fragments (with two unique anchors)
umDT <- rbindlist(lapply(names(BPEL), function(s){
  agr <- BPEL[[s]]
  uniq.gr <- agr[mcols(agr)[["NH1"]] %in% 1 | mcols(agr)[["NH2"]] %in% 1]
  rdt <- data.table("sample" = s,
                    "input_seq" = length(agr),
                    "unique_anchor_seq" = length(uniq.gr))
  rdt[,"percent_unique" := round((unique_anchor_seq/input_seq)*100,2)]
  return(rdt)})); umDT

## Calculate library size factor - use the total input
umDT[,"min_input" := min(input_seq)]
umDT[,"szf" := input_seq/min_input]
umDT

##
## CALCULATE STRANDED COVERAGE
##

## - NOTE: don't use reads since we do not have UMIs
## - NOTE: only use perfect mappers

coverage.SL <- lapply(names(BPEL), function(s){
  message(paste0("sample: ",s))
  agr <- BPEL[[s]]
  ngr <- agr[mcols(agr)[["NH1"]] %in% 1 | mcols(agr)[["NH2"]] %in% 1]
  total_sequences <- length(ngr)

  message("\tcoverage by strand")
  plus.cov <- coverage(ngr[strand(ngr) %in% "+"])
  minus.cov <- coverage(ngr[strand(ngr) %in% "-"])

  message("\tsaving a list")
  coverage.list <- list("watson"=plus.cov,"crick"=minus.cov)
```

```
  return(coverage.list)})
names(coverage.SL) <- names(BPEL)

## save image
rm(BPEL,bac_order)
save.image(file = "/Users/genzorp/Documents/GITHUB/LIVE/piggyBac_mutants/data/PB_peakNcov.RData")
```

### PLOTTING RESULTS

Plot chromosome peak distribution between various samples.

```
## NOTE: Load previous object
rm(list = ls())
load(file = "/Users/genzorp/Documents/GITHUB/LIVE/piggyBac_mutants/data/PB_peakNcov.RData")

## library
library(scales)
```

```
##
## Attaching package: 'scales'

## The following object is masked from 'package:purrr':
##
##     discard

## The following object is masked from 'package:readr':
##
##     col_factor
```

```
## View info about data
dtsi
```

```
##                               original    te1    te2       genotype    rep
##                                 <char> <char> <char>         <char> <char>
##  1: HCT116_le40_le40_delta74hyPB2CD_1   le40   le40 delta74hyPB2CD      1
##  2: HCT116_le40_le40_delta74hyPB2CD_2   le40   le40 delta74hyPB2CD      2
##  3: HCT116_le40_le40_delta74hyPB2CD_3   le40   le40 delta74hyPB2CD      3
##  4:   HCT116_le40_le40_delta74PB2CD_1   le40   le40   delta74PB2CD      1
##  5:   HCT116_le40_le40_delta74PB2CD_2   le40   le40   delta74PB2CD      2
##  6:   HCT116_le40_le40_delta74PB2CD_3   le40   le40   delta74PB2CD      3
##  7:           HCT116_re67_le40_hyPB_1   re67   le40           hyPB      1
##  8:           HCT116_re67_le40_hyPB_2   re67   le40           hyPB      2
##  9:           HCT116_re67_le40_hyPB_3   re67   le40           hyPB      3
## 10:             HCT116_re67_le40_PB_1   re67   le40             PB      1
## 11:             HCT116_re67_le40_PB_2   re67   le40             PB      2
## 12:             HCT116_re67_le40_PB_3   re67   le40             PB      3
##     peak_count
##          <int>
##  1:       3998
```

```
##  2:       3844
##  3:       4648
##  4:       3235
##  5:       2846
##  6:       3172
##  7:       7626
##  8:       7058
##  9:       7676
## 10:       9298
## 11:       9107
## 12:       9311
```

```r
## Create objects governing the order
all_order
```

```
##  [1] "HCT116_re67_le40_PB_1"              "HCT116_re67_le40_PB_2"
##  [3] "HCT116_re67_le40_PB_3"              "HCT116_re67_le40_hyPB_1"
##  [5] "HCT116_re67_le40_hyPB_2"            "HCT116_re67_le40_hyPB_3"
##  [7] "HCT116_le40_le40_delta74PB2CD_1"   "HCT116_le40_le40_delta74PB2CD_2"
##  [9] "HCT116_le40_le40_delta74PB2CD_3"   "HCT116_le40_le40_delta74hyPB2CD_1"
## [11] "HCT116_le40_le40_delta74hyPB2CD_2" "HCT116_le40_le40_delta74hyPB2CD_3"
```

```r
group_order <- unique(unlist(lapply(all_order,function(s){substr(x = s, start = 1,
                                                                 stop = nchar(s)-2)})))
group_cols <- c("#333333","#0096B8","#F85A3A","#E09D00")

## peak count by chromosome
chr_pcnt <- rbindlist(lapply(names(peak.SL),function(s){
  pgr <- peak.SL[[s]]
  adt <- as.data.table(pgr)
  cdt <- adt[,.N,by=c("sample","seqnames")]
  return(cdt) })); chr_pcnt
```

```
##                                 sample seqnames     N
##                                 <char>   <fctr> <int>
##   1: HCT116_le40_le40_delta74hyPB2CD_1     chr1   347
##   2: HCT116_le40_le40_delta74hyPB2CD_1     chr2   232
##   3: HCT116_le40_le40_delta74hyPB2CD_1     chr3   199
##   4: HCT116_le40_le40_delta74hyPB2CD_1     chr4   291
##   5: HCT116_le40_le40_delta74hyPB2CD_1     chr5   161
##  ---
## 272:             HCT116_re67_le40_PB_3    chr19   267
## 273:             HCT116_re67_le40_PB_3    chr20   263
## 274:             HCT116_re67_le40_PB_3    chr21    52
## 275:             HCT116_re67_le40_PB_3    chr22   177
## 276:             HCT116_re67_le40_PB_3     chrX   135
```

```r
## Add group and total per group
chr_pcnt[,"group" := substring(sample, 1, nchar(sample)-2)]
colnames(chr_pcnt) <- gsub("N","count",colnames(chr_pcnt))
chr_pcnt[,"total" := sum(.SD), by = "group", .SDcols = "count"]
chr_pcnt[,"percent" := round((count/total)*100,3)]
```

```r
## Calculate mean percent per chromosome
chrc.se <- setDT(summarySE(data = chr_pcnt, groupvars = c("group","seqnames"),
                           measurevar = "percent"))
chrc.se[["group"]] <- factor(chrc.se[["group"]], levels = group_order)

## Order the group
chrc.se[["group"]] <- factor(chrc.se[["group"]], levels = rev(group_order))

## Plot heatmap
genTile <- ggplot() + theme_pubclean() +
  geom_tile(data = chrc.se, aes(y = group, x = seqnames, fill=percent),
            colour = "white", size = 1) +
  scale_fill_distiller(palette = "PuBu", direction = 1) +
  coord_fixed() + xlab("") + ylab("") +
  theme(axis.ticks = element_blank(),
        panel.grid.minor.x = element_blank(),panel.grid.minor.y = element_blank(),
        panel.grid.major.x = element_blank(),panel.grid.major.y = element_blank(),
        axis.text.x = element_text(angle = 90, hjust = 1, vjust = 0.5))

## Plot
genTile
```

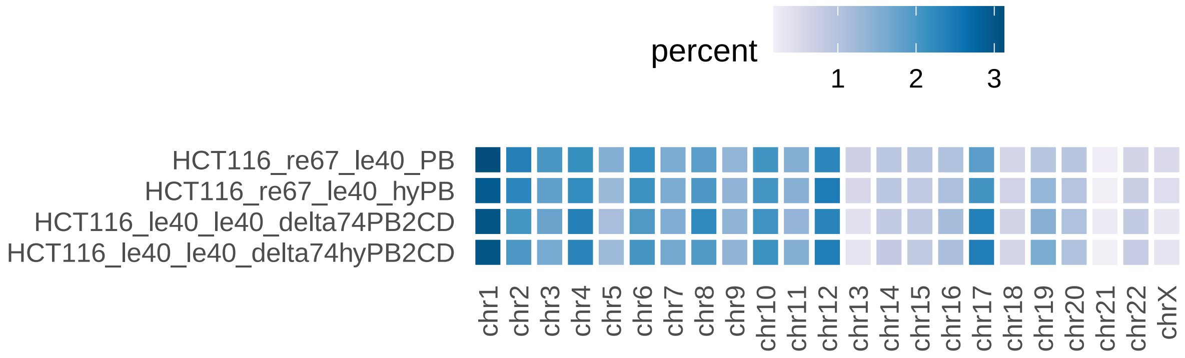


#### Genome annotation

Plot the genomic distribution of various annotation categories for relative comparison.

```r
## NOTE: The specific curated genome annotation table used here is available upon request

## Load curated human genome annotation
load(file = "/Users/genzorp/Documents/GITHUB/PG_Annotations/sessions/gencodeV38_hg38_curated.RData")

## Create genome-specific information table
hg38si <- as.data.table(keepStandardChromosomes(seqinfo(Hsapiens)), keep.rownames=TRUE)
```

```
## Warning in as.data.frame.Seqinfo(x, ...): extra arguments were ignored
```

```r
hg38si[["start"]] <- 1
colnames(hg38si) <- gsub("rn","seqnames",colnames(hg38si))
colnames(hg38si) <- gsub("seqlengths","end",colnames(hg38si))
hg38si[["seqnames"]] <- factor(hg38si[["seqnames"]], levels = hg38si[["seqnames"]])

## annotation GR
pc.GENIC <- reduce(gencode38[mcols(gencode38)[["gene_type"]] %in% "protein_coding"],
                   ignore.strand=TRUE)
all.GENIC <- reduce(gencode38, ignore.strand=TRUE)
all.INTERGENIC <- gaps(all.GENIC)

## category genome proportions
genomeSize <- sum(hg38si[["end"]])
gf.GENIC <- sum(width(all.GENIC))
gf.INTERGENIC <- sum(width(all.INTERGENIC))
gf.pc.GENIC <- sum(width(pc.GENIC))
gf.pc.EXONS <- sum(width(reduce(exons_gc38,ignore.strand=TRUE)))
gf.pc.INTRONS <- sum(width(reduce(introns_gc38,ignore.strand=TRUE)))

## category fractions
fr_GENIC <- (gf.GENIC/genomeSize)*100
fr_INTERGENIC <- 100 - fr_GENIC
fr_pcGENIC <- (gf.pc.GENIC/genomeSize)*100
fr_EXONS <- (gf.pc.EXONS/genomeSize)*100
fr_INTRONS <- fr_pcGENIC-fr_EXONS

##
## CREATE SEPARATE PLOTS
##

## GENIC vs. INTERGENIC

## table
GDT_A <- data.table("fragment"= factor(x = c("genic","intergenic")),
                    "fraction"=c(fr_GENIC,fr_INTERGENIC)); GDT_A
```

```
##      fragment fraction
##        <fctr>    <num>
## 1:      genic   56.876
## 2: intergenic   43.124
```

```r
## plot
pie_A <- ggplot(GDT_A,aes(x = "", y = fraction, fill = fct_inorder(fragment))) +
  geom_col(width = 1, color = "white" , size = 2) +
  geom_text(aes(label = round(fraction,0)),
            position = position_stack(vjust = 0.5), size = 6, colour = "white") +
  ggtitle("fraction of GENOME (%)") +
  coord_polar(theta = "y",start = 0) +
  scale_fill_brewer(palette = "Set1") + theme_void()

## PROTEIN CODING vs. NON CODING

## table
```

```r
GDT_B <- data.table("fragment"= factor(x = c("protein_coding","non_coding")),
                    "fraction"=c((fr_pcGENIC/fr_GENIC)*100,
                                 100-((fr_pcGENIC/fr_GENIC)*100))); GDT_B
```

```
##          fragment fraction
##            <fctr>    <num>
## 1: protein_coding 73.85608
## 2:     non_coding 26.14392
```

```r
## plot
pie_B <- ggplot(GDT_B,aes(x = "", y = fraction, fill = fct_inorder(fragment))) +
  geom_col(width = 1, color = "white" , size = 2) +
  geom_text(aes(label = round(fraction,0)),
            position = position_stack(vjust = 0.5), size = 6, colour = "white") +
  ggtitle("fraction of GENIC space (%)") +
  coord_polar(theta = "y",start = 0) +
  scale_fill_brewer(palette = "Set1") + theme_void()

## PROTEIN CODING EXONS vs INTRONS

## table
GDT_C <- data.table("fragment"= factor(x = c("protein_coding_exons","protein_coding_introns")),
                    "fraction"=c((fr_EXONS/fr_pcGENIC)*100,
                                 100-((fr_EXONS/fr_pcGENIC)*100))); GDT_C
```

```
##                  fragment  fraction
##                    <fctr>     <num>
## 1:   protein_coding_exons  8.092603
## 2: protein_coding_introns 91.907397
```

```r
## plot
pie_C <- ggplot(GDT_C,aes(x = "", y = fraction, fill = fct_inorder(fragment))) +
  geom_col(width = 1, color = "white" , size = 2) +
  geom_text(aes(label = round(fraction,0)),
            position = position_stack(vjust = 0.5), size = 6, colour = "white") +
  ggtitle("fraction of PROTEIN CODING genes (%)") +
  coord_polar(theta = "y",start = 0) +
  scale_fill_brewer(palette = "Set1") + theme_void()

## Combine all plots into a figure
annotPies <- ggarrange(plotlist = list(pie_A, pie_B, pie_C), ncol = 1, nrow = 3)
annotPies
```

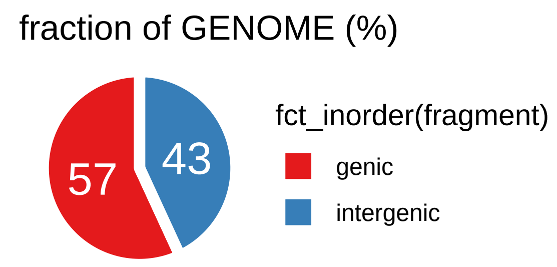


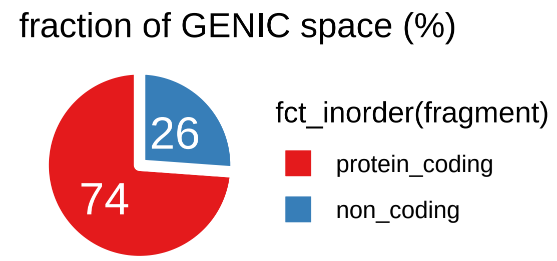


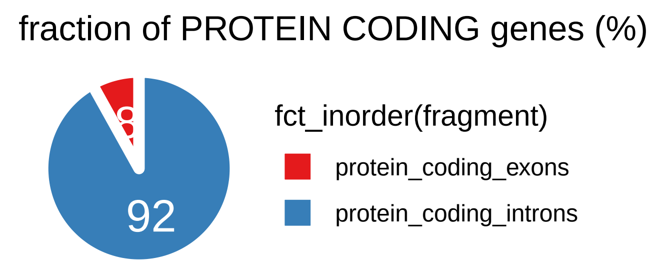


Calculate the overlaps of the peaks with different genome features.

```r
## Plot specific
FAM = "Helvetica"; XYT = 8; TCOL="black"
group_cols <- c("#333333", "#0096B8", "#F85A3A", "#E09D00")

## Data
load(file = "/Users/genzorp/Documents/GITHUB/LIVE/piggyBac_mutants/data/PB_peakNcov.RData")
load(file = "/Users/genzorp/Documents/GITHUB/PG_Annotations/sessions/gencodeV38_hg38_curated.RData")
#peak.SL;gencode38

## annotation GR
pc.GENIC <- reduce(gencode38[mcols(gencode38)[["gene_type"]] %in% "protein_coding"],
                   ignore.strand=TRUE)
all.GENIC <- reduce(gencode38, ignore.strand=TRUE)
all.INTERGENIC <- gaps(all.GENIC)

## category genome proportions
genomeSize <- sum(hg38si[["end"]])
gf.GENIC <- sum(width(all.GENIC))
gf.INTERGENIC <- sum(width(all.INTERGENIC))
gf.pc.GENIC <- sum(width(pc.GENIC))
gf.pc.EXONS <- sum(width(reduce(exons_gc38,ignore.strand=TRUE)))
gf.pc.INTRONS <- sum(width(reduce(introns_gc38,ignore.strand=TRUE)))

## category fractions
fr_GENIC <- (gf.GENIC/genomeSize)*100
```

```r
fr_INTERGENIC <- 100 - fr_GENIC
fr_pcGENIC <- (gf.pc.GENIC/genomeSize)*100
fr_EXONS <- (gf.pc.EXONS/genomeSize)*100
fr_INTRONS <- fr_pcGENIC-fr_EXONS

## groups
all.GENIC <- reduce(gencode38, ignore.strand=TRUE)
all.INTERGENIC <- gaps(all.GENIC)

##
## GENIC
##

## fraction of peaks in genes
all.GENIC.dt <- rbindlist(lapply(names(peak.SL), function(s){
  apgr <- peak.SL[[s]]
  pgr <- apgr[mcols(apgr)[["peak_height"]] >= 5]
  inFEAT <- subsetByOverlaps(x = pgr, ranges = all.GENIC)
  gDT <- data.table("sample"=s,
                    "total_peaks"=length(pgr),
                    "peaks_inFEAT"=length(inFEAT),
                    "fraction_inFEAT"=(length(inFEAT)/length(pgr)*100))
  gDT[["feature"]] <- "all_genic"
  return(gDT) }) )

## summarize & organize
all.GENIC.dt[,"group":= substr(sample,1,(nchar(sample)-2))]
genic_count_dtse <- summarySE(data = all.GENIC.dt,
                              groupvars = "group",
                              measurevar = "peaks_inFEAT")
genic_dtse <- summarySE(data = all.GENIC.dt,
                        groupvars = "group",
                        measurevar = "fraction_inFEAT")

genic_dtse[["group"]] <- factor(genic_dtse[["group"]],levels = rev(group_order))

## calculate change
genic_dtse <- setDT(genic_dtse)
genic_dtse[,"feat" := fr_GENIC]
genic_dtse[,"diff" := fraction_inFEAT - feat]
genic_dtse[,"annotation" := "genic_intergenic"]

##
## PROTEIN CODING vs NON-CODING
##

## groups
pc.GENIC <- reduce(gencode38[mcols(gencode38)[["gene_type"]] %in% "protein_coding"],
                   ignore.strand=TRUE)
pc.NC <- reduce(gencode38[!mcols(gencode38)[["gene_type"]] %in% "protein_coding"],
                ignore.strand=TRUE)

## fraction of peaks in protein coding genes
```

```
sub.GENIC.dt <- rbindlist(lapply(names(peak.SL), function(s){
  apgr <- peak.SL[[s]]
  pgr <- apgr[mcols(apgr)[["peak_height"]] >= 5]
  inFEAT_A <- subsetByOverlaps(x = pgr, ranges = pc.GENIC)
  inFEAT_B <- subsetByOverlaps(x = pgr, ranges = pc.NC)
  gDT <- data.table("sample"=s,
                    "total_peaks"=length(pgr),
                    "peaks_inFEAT_A"=length(inFEAT_A),
                    "peaks_inFEAT_B"=length(inFEAT_B))
  gDT[["feature_A"]] <- "pc_genic"
  gDT[["feature_B"]] <- "pc_nc"
  return(gDT) }) )

## Summarize
sub.GENIC.dt[,"featuresTotal" := peaks_inFEAT_A+peaks_inFEAT_B]
sub.GENIC.dt[,"fraction_inFEAT" := (peaks_inFEAT_A/featuresTotal)*100]

## summarize & organize
sub.GENIC.dt[,"group":= substr(sample,1,(nchar(sample)-2))]
sub.GENIC_count_dtse <- summarySE(data = sub.GENIC.dt,
                  groupvars = "group", measurevar = "peaks_inFEAT_A")
sub.GENIC_dtse <- summarySE(data = sub.GENIC.dt,
                  groupvars = "group", measurevar = "fraction_inFEAT")
sub.GENIC_dtse[["group"]] <- factor(sub.GENIC_dtse[["group"]],levels = group_order)


## calculate change
sub.GENIC_dtse <- setDT(sub.GENIC_dtse)
sub.GENIC_dtse[,"feat" := (fr_pcGENIC/fr_GENIC)*100]
sub.GENIC_dtse[,"diff" := fraction_inFEAT - feat]
sub.GENIC_dtse[,"annotation" := "protCoding_nonCoding"]

##
## INTRONS vs EXONS
##

## groups
pc.EXONS <-reduce(exons_gc38,ignore.strand=TRUE)
pc.INTRONS <- reduce(introns_gc38,ignore.strand=TRUE)

## fraction of peaks inexons
in.EXON.dt <- rbindlist(lapply(names(peak.SL), function(s){
  apgr <- peak.SL[[s]]
  pgr <- apgr[mcols(apgr)[["peak_height"]] >= 5]
  inFEAT_A <- subsetByOverlaps(x = pgr, ranges = pc.EXONS)
  inFEAT_B <- subsetByOverlaps(x = pgr, ranges = pc.INTRONS)
  gDT <- data.table("sample"=s,
                    "total_peaks"=length(pgr),
                    "peaks_inFEAT_A"=length(inFEAT_A),
                    "peaks_inFEAT_B"=length(inFEAT_B))
  gDT[["feature_A"]] <- "pc_exon"
  gDT[["feature_B"]] <- "pc_intron"
  return(gDT) }) )
```

```r
## Summarize
in.EXON.dt[,"featuresTotal" := peaks_inFEAT_A+peaks_inFEAT_B]
in.EXON.dt[,"fraction_inFEAT" := (peaks_inFEAT_A/featuresTotal)*100]

## summarize & organize
in.EXON.dt[,"group":= substr(sample,1,(nchar(sample)-2))]
in.EXON_count_dtse <- summarySE(data = in.EXON.dt,
                  groupvars = "group", measurevar = "peaks_inFEAT_A")
in.EXON_dtse <- summarySE(data = in.EXON.dt,
                  groupvars = "group", measurevar = "fraction_inFEAT")
in.EXON_dtse[["group"]] <- factor(in.EXON_dtse[["group"]],levels = group_order)

## calculate change
in.EXON_dtse <- setDT(in.EXON_dtse)
in.EXON_dtse[,"feat" := (fr_EXONS/fr_pcGENIC)*100]
in.EXON_dtse[,"diff" := fraction_inFEAT - feat]
in.EXON_dtse[,"annotation" := "exons_introns"]

## Combine tables
ann.dt <- rbindlist(l = list(genic_dtse,sub.GENIC_dtse,in.EXON_dtse))

##
## PLOT DIRECT
##

## organize
ann.dt[["group"]] <- factor(ann.dt[["group"]],levels = group_order)
ann.dt[["annotation"]] <- factor(ann.dt[["annotation"]],
                                 levels = c("genic_intergenic","protCoding_nonCoding",
                                            "exons_introns"))

## Plot
barAnnotUp <- ggplot() + theme_pubclean() +
  geom_bar(data = ann.dt,
                   aes(x = annotation, y = fraction_inFEAT, fill = group, group = group),
                   stat = "identity", position = "dodge", colour = "white",
                   width = 0.75) +
  #geom_hline(yintercept = unique(ann.dt[["feat"]]), linetype="dashed", colour = "firebrick") +
  geom_errorbar(data = ann.dt,
                aes(x = annotation, ymin = fraction_inFEAT - se, ymax = fraction_inFEAT + se,
                   group = group),colour = "black",
                position = position_dodge(width = 0.75),
                width = 0.4) +
  geom_text(data = ann.dt,
            aes(x = annotation, y = fraction_inFEAT+3, label = round(fraction_inFEAT,0),
                group = group),
            position = position_dodge(width = 0.75)) +
  ylab("% peaks in the annotaion group") +
  scale_colour_manual(values = group_cols) +
  scale_fill_manual(values = group_cols) +
  theme(aspect.ratio = 0.5, legend.position = "bottom",
        axis.text.y = element_text(family = FAM, size = XYT, colour = TCOL))
```

```
## Plot
barAnnotUp
```

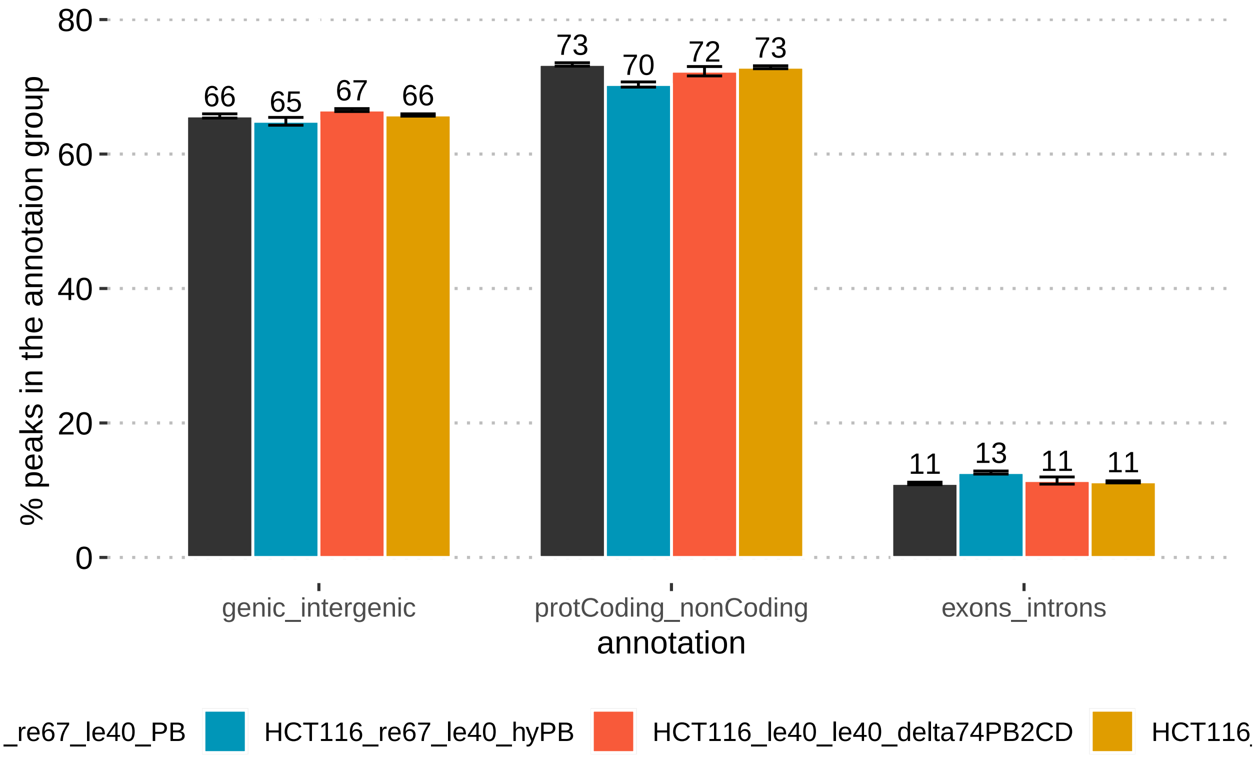


#### Tiled integration map

The genome has been split into 1 Mb large tiles, an various features within each tile were compared.

```
## NOTE: Load previous object
rm(list = ls())
load(file = "/Users/genzorp/Documents/GITHUB/LIVE/piggyBac_mutants/data/PB_peakNcov.RData")

##
## SPLIT GENOME INTO 1MB TILES
##

## Create a genome-wide tiles
hg38.tile <- tileGenome(seqlengths = keepStandardChromosomes(seqinfo(Hsapiens)),
                        tilewidth = 1000000, cut.last.tile.in.chrom = TRUE)
hg38.tdt <- as.data.table(hg38.tile)
hg38.tdt[,"tileN" := paste0(seqnames,"_",1:nrow(.SD)), by=seqnames]
hg38.tgr <- makeGRangesFromDataFrame(df = hg38.tdt, keep.extra.columns = TRUE)
names(hg38.tgr) <- mcols(hg38.tgr)[["tileN"]]

##
## CENTROMERES
##
```

```r
## Load public centromere information
centro.path <- "/Users/genzorp/Documents/GITHUB/PG_Annotations/data/Hsapiens/centromeres_hg38/cytobandi
centro.gr <- makeGRangesFromDataFrame(fread(centro.path),keep.extra.columns = TRUE)
centro.grl <- split(centro.gr, ~seqnames)
centro.fgr <- unlist(range(centro.grl))
centro.fdt <- as.data.table(centro.fgr)

## Make centromere table
hg38.centro.dt <- as.data.table(
hg38.tgr[queryHits(findOverlaps(query = hg38.tgr, subject = centro.fgr))])
hg38.centro.dt[["tileN"]] <- as.integer(unlist(tstrsplit(hg38.centro.dt[["tileN"]],
                                                          split="_",keep = 2)))
hg38.centro.dt <- setDT(hg38.centro.dt)
hg38.centro.dt[,"category" := "centrosome"]
hg38.centro.dt[,"value" := 1]
hg38.centro.dt[,"rescaled" := 1]
hg38.centro.dt <- hg38.centro.dt[,.SD,.SDcols=c("seqnames","tileN","category","value","rescaled")]

##
## NUCLEOTIDE FREQUENCIES
##

## Extract genomic sequence for each tile (takes little time)
hg38.tgr.seq <- getSeq(Hsapiens, hg38.tgr)

## Calculate DINUCLEOTIDE frequency per tile
hg38.di.tgr <- hg38.tgr
mcols(hg38.di.tgr) <- dinucleotideFrequency(x = hg38.tgr.seq, as.prob = TRUE)
hg38.di.dt <- as.data.table(x = hg38.di.tgr)
hg38.di.dt[["names"]] <- names(hg38.di.tgr)
hg38.di.dt <- setDT(hg38.di.dt)
hg38.di.dt[,"tileN" := tstrsplit(names,split="_",keep=2)]
hg38.di.dt[["tileN"]] <- as.numeric(hg38.di.dt[["tileN"]])

## Melt table to extract desired frequencies
hg38.di.dtm  <- melt.data.table(data = hg38.di.dt, variable.name = "category",
                                value.name = "value",
                                id.vars = c("seqnames","tileN"),
                                measure.vars = c("GC"))
hg38.di.dtm[["rescaled"]] <- scales::rescale(x = hg38.di.dtm[["value"]], to = c(0,1))

## Calculate TETRANUCLEOTIDE frequency per tile
hg38.tetra.tgr <- hg38.tgr
mcols(hg38.tetra.tgr) <- oligonucleotideFrequency(x = hg38.tgr.seq, as.prob = TRUE, width = 4)
hg38.tetra.dt <- as.data.table(x = hg38.tetra.tgr)
hg38.tetra.dt[["names"]] <- names(hg38.tetra.tgr)
hg38.tetra.dt <- setDT(hg38.tetra.dt)
hg38.tetra.dt[,"tileN" := tstrsplit(names,split="_",keep=2)]
hg38.tetra.dt[["tileN"]] <- as.numeric(hg38.tetra.dt[["tileN"]])

## Melt table to extract desired frequencies
hg38.tetra.dtm  <- melt.data.table(data = hg38.tetra.dt, variable.name = "category",
                                   value.name = "value",
```

```
                                   id.vars = c("seqnames","tileN"),
                                   measure.vars = c("TTAA"))
hg38.tetra.dtm[["rescaled"]] <- scales::rescale(x = hg38.tetra.dtm[["value"]], to = c(0,1))

##
## GENE ABUNDANCE
##

load(file = "/Users/genzorp/Documents/GITHUB/PG_Annotations/sessions/gencodeV38_hg38_curated.RData")
hg38.genes.tgr <- hg38.tgr
mcols(hg38.genes.tgr)[["value"]] <- countOverlaps(query = hg38.genes.tgr, subject = gencode38)
hg38.genes.dt <- as.data.table(x = hg38.genes.tgr)
hg38.genes.dt[["names"]] <- names(hg38.genes.tgr)
hg38.genes.dt[,"tileN" := tstrsplit(names,split="_",keep=2)]
hg38.genes.dt[["tileN"]] <- as.numeric(hg38.genes.dt[["tileN"]])
hg38.genes.dt[["category"]] <- factor(x = "genes")
hg38.genes.dt <- hg38.genes.dt[,.SD,.SDcols = c("seqnames","tileN","category","value")]
hg38.genes.dt[["rescaled"]] <- scales::rescale(x = hg38.genes.dt[["value"]], to = c(0,1))

##
## PIGGYBAC PEAKS
##

## Count number of peaks per tile
hg38.tgr.peaks.dt <- rbindlist(lapply(names(peak.SL), function(s){
  pgr <- peak.SL[[s]]
  mcols(hg38.tile)[["count"]] <- countOverlaps(query = hg38.tgr,
                                              subject = pgr,
                                              ignore.strand=TRUE)
  adt <- as.data.table(hg38.tile)
  adt[["sample"]] <- s
  adt <- setDT(adt)
  adt[,"tileN" := 1:nrow(.SD), by = seqnames]
  return(adt)}))

## Normalize by calculating fraction of total peaks per sample
hg38.tgr.peaks.dt[,"sample_total" := sum(.SD), by="sample", .SDcols="count"]
hg38.tgr.peaks.dt[,"value" := round((count/sample_total)*100,5)]

## summarize replicates and organize: mean percentage of peaks per tile
hg38.tgr.peaks.dt[,"category" := substring(sample,1,nchar(sample)-2)]
hg38.tgr.peaks.se <- setDT(summarySE(data = hg38.tgr.peaks.dt,
                                     groupvars = c("seqnames","tileN","category"),
                                     measurevar = "value"))
group_order <- c("HCT116_re67_le40_PB","HCT116_re67_le40_hyPB",
                 "HCT116_le40_le40_delta74PB2CD","HCT116_le40_le40_delta74hyPB2CD")
hg38.tgr.peaks.se[["category"]] <- factor(hg38.tgr.peaks.se[["category"]],
                                          levels = rev(group_order))
hg38.tgr.peaks.se[["rescaled"]] <- scales::rescale(x = hg38.tgr.peaks.se[["value"]],
                                                   to = c(0,1))
hg38.tgr.peaks.se <- hg38.tgr.peaks.se[,.SD,.SDcols=c("seqnames","tileN",
                                                      "category","value","rescaled")]
```

```r
##
## COMBINE DATA
##

hg38.gt <- rbindlist(l = list(hg38.tgr.peaks.se,hg38.centro.dt,
                              hg38.tetra.dtm,hg38.di.dtm,hg38.genes.dt))
hg38.gt[["category"]] <- factor(hg38.gt[["category"]],
                                levels = rev(c(group_order,"centrosome",
                                               "TTAA","GC","genes")))
hg38.gt[["tileN"]] <- as.numeric(hg38.gt[["tileN"]] )
hg38.gt <- setDT(hg38.gt)
hg38.gt[,"rescaled2" := ifelse(rescaled == 0, NA, rescaled)]

##
## MAKE THE PLOT
##

genomicFEATURES <- ggplot() + theme_void() +
  geom_tile(data = hg38.gt,
            aes(x = tileN, y = category, fill = rescaled2), colour = "white") +
  scale_fill_distiller(palette = "Spectral", na.value = "#DEE5E5",direction = -1,
                       name="Normalized\nfraction") +
  facet_wrap(~seqnames, ncol = 1) + ylab("") +
  ggtitle(paste0("Tiled map of genome (each tile is 1 Mb)\n",
                 "Row order = ",paste0(group_order, collapse = ", "),
                 "\ncentrosome ",",",TTAA ",",",GC ",",",genes")) +
  theme(aspect.ratio = 0.035, legend.position = "right",
        strip.text = element_text(hjust = 0),
        axis.text.y = element_blank());

## Print the plot
genomicFEATURES
```

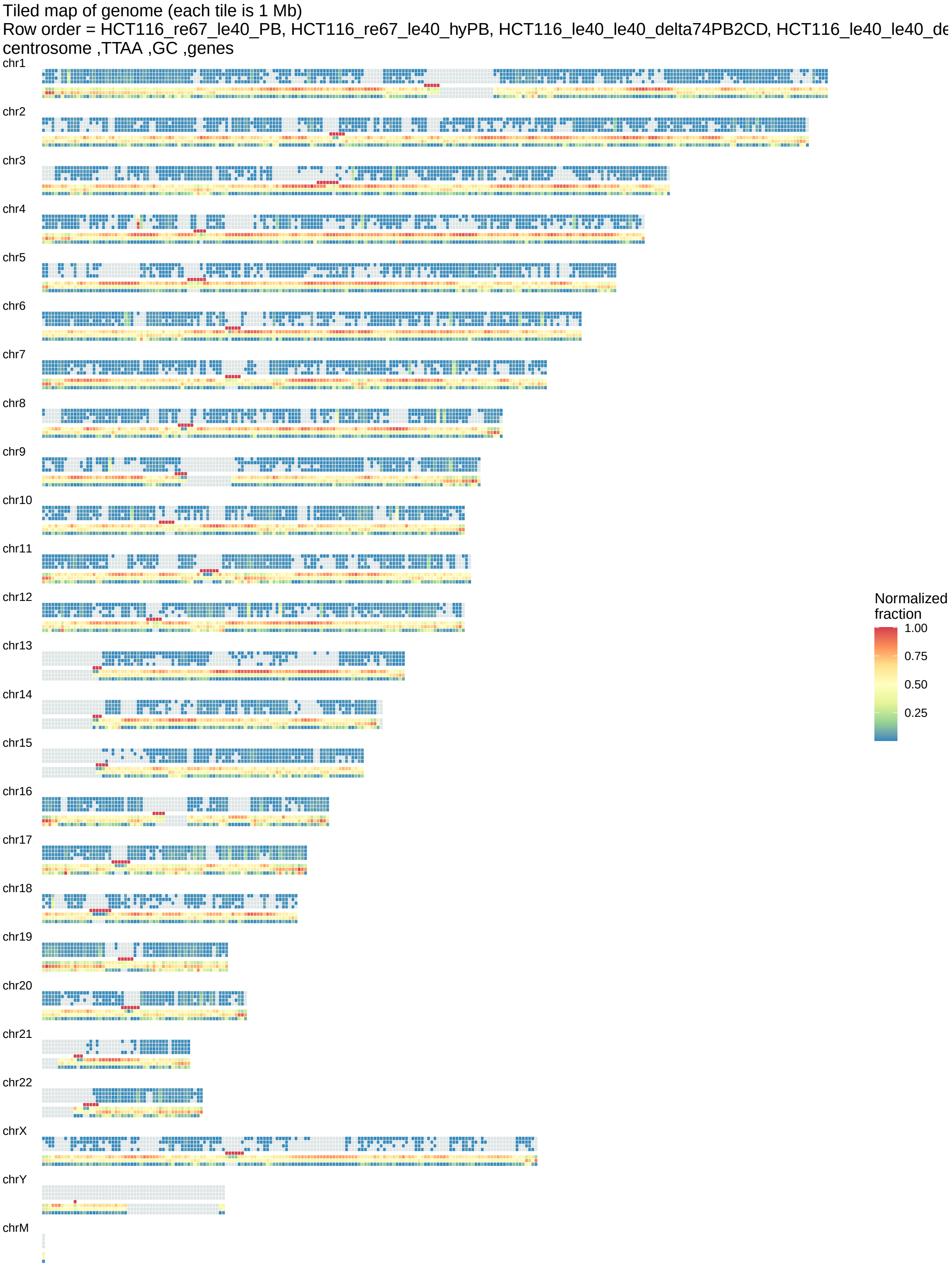
Tiled map of genome (each tile is 1 Mb)
Row order = HCT116_re67_le40_PB, HCT116_re67_le40_hyPB, HCT116_le40_le40_delta74PB2CD, HCT116_le40_le40_de
centrosome ,TTAA ,GC ,genes
chr1
chr2
chr3
chr4
chr5
chr6
chr7
chr8
chr9
chr10
chr11
chr12
chr13
chr14
chr15
chr16
chr17
chr18
chr19
chr20
chr21
chr22
chrX
chrY
chrM
Normalized fraction
1.00
0.75
0.50
0.25

**Genome coverage**

```r
## NOTE: previous chunk is REQUIRED
group_cols <- c("#333333","#0096B8","#F85A3A","#E09D00")

## Select a tile from browsing a map
chr7t131.dt <- hg38.tgr.peaks.se[seqnames %in% "chr7"][tileN %in% 131]
chr7t131.gr <- hg38.tgr[mcols(hg38.tgr)[["tileN"]] %in% "chr7_131"]
chr7t131.gr.org <- chr7t131.gr

## change coordinates
new_start = 130100000
new_end =   131600000

## Change coordinates
start(chr7t131.gr) <- new_start
end(chr7t131.gr) <- new_end

##
## Setup sliding window
##

## Sliding window for extended tile window
chr7t131.sw <- unlist(slidingWindows(x = chr7t131.gr, width = 5000, step = 500))

##
## Calculate k-mer frequencies
##

## TTAA
chr7t131.sw.seq <- getSeq(Hsapiens,chr7t131.sw)
mcols(chr7t131.sw)[["TTAA"]] <- oligonucleotideFrequency(x = chr7t131.sw.seq,
                                                         width = 4, as.prob = TRUE)[,"TTAA"]
ttaa.freq.dt <- as.data.table(chr7t131.sw)
ttaa.freq.dt[,"mid" := start+(width/2)]
ttaa.freq.dt[["sample"]] <- "C_TTAA_content"

## GC
mcols(chr7t131.sw)[["GC"]] <- dinucleotideFrequency(x = chr7t131.sw.seq, as.prob = TRUE)[,"GC"]
gc.freq.dt <- as.data.table(chr7t131.sw)
gc.freq.dt[,"mid" := start+(width/2)]
gc.freq.dt[["sample"]] <- "D_GC_content"

##
## Location of the genes
##

## Number of genes
genes.in.tile <- subsetByOverlaps(x = gencode38, ranges = chr7t131.gr)
```

```
genes.in.tile.grl <- split(genes.in.tile,~gene_id)
genes.in.tile.gr <- unlist(GenomicRanges::reduce(genes.in.tile.grl))
mcols(genes.in.tile.gr)[["gene_id"]] <- names(genes.in.tile.gr)
genes.in.tile.dt <- as.data.table(genes.in.tile.gr)
genes.in.tile.dt[["sample"]] <- factor(x = c("G_genes"))
genes.in.tile.dt[["y"]] <- seq(0.1,0.2*nrow(genes.in.tile.dt),0.2)

##
## Read coverage of peaks
##

## ensure same leves and names are used
seqlevels(chr7t131.sw) <- seqlevelsInUse(chr7t131.sw)

## Get normalized coverage per sample
coverage.dt <- suppressWarnings(
  rbindlist(lapply(names(coverage.SL), function(s){
  stranded.cov <- coverage.SL[[s]]
  unstranded.cov <- stranded.cov$watson + stranded.cov$crick
  usedSeqLevel <- seqlevels(chr7t131.sw)
  unstranded.cov <- unstranded.cov[names(unstranded.cov) %in% usedSeqLevel]
  lib_size <- umDT[sample %in% s][["input_seq"]]
  #message(paste0("sample: ",s,"\nlibrary size: ", lib_size))
  norm.unstranded.cov <- unstranded.cov/(lib_size/1000000)
  coverage.gr <- binnedAverage(bins = chr7t131.sw, numvar = norm.unstranded.cov, varname = "ncov")
  coverage.dt <- as.data.table(coverage.gr)
  coverage.dt[["full_sample"]] <- s
  return(coverage.dt)})))

## Calculate mean normalized coverage per group
coverage.dt[,"group" := substr(full_sample, 1, nchar(full_sample)-2)]
coverage.dtm <- setDT(summarySE(data = coverage.dt,
                                groupvars = c("seqnames","start","end","group"),
                                measurevar = c("ncov")))
coverage.dtm[,"mid":= start+((end-start)/2)]
coverage.dtm[["group"]] <- factor(coverage.dtm[["group"]], levels = group_order)

##
## piggyBac peaks
##

peak_extention = 1000
pb.peak.dt <- rbindlist(lapply(group_order, function(g){
  agrl <- peak.SL[grep(g,names(peak.SL))]
  cgr <- c(agrl[[1]],agrl[[2]],agrl[[3]])
  cgr <- subsetByOverlaps(x = cgr, ranges = chr7t131.sw)
  start(cgr) <- start(cgr) - peak_extention
  end(cgr) <- end(cgr) + peak_extention
  cgrr <- GenomicRanges::reduce(cgr,ignore.strand=TRUE, with.revmap=TRUE)
  cgrr.dt <- as.data.table(cgrr)
  cgrr.dt[["group"]] <- g
  cgrr.dt[["revmap"]] <- NULL
  return(cgrr.dt) }) )
```

```r
pb.peak.dt[["group"]] <- factor(pb.peak.dt[["group"]], levels = group_order)

## Plotting objects
gc.freq.dt[["group"]] <- "A_GC"
gc.freq.dt[["group"]] <- factor(gc.freq.dt[["group"]], levels = c("A_GC"))
ttaa.freq.dt[["group"]] <- "B_TTAA"
ttaa.freq.dt[["group"]] <- factor(ttaa.freq.dt[["group"]],levels = c("B_TTAA"))
genes.in.tile.dt[["group"]] <- "C_GENES"
genes.in.tile.dt[["group"]] <- factor(genes.in.tile.dt[["group"]],levels = c("C_GENES"))

## Scale constant
aK = 100

## PB
pcA <- ggplot() + theme_pubclean() +
  ## coverage
  geom_area(data = coverage.dtm,
            aes(x = mid, y = ncov, fill = group),
            size = 0.3, colour = "black") +
  ## peaks
  geom_segment(data = pb.peak.dt,
               aes(x = start, xend = end, y=10, yend = 10, colour = group),
               size = 6) +
  ## GC
  geom_step(data = gc.freq.dt, aes(x = mid, y = GC*aK),
            size = 0.3, colour = "black") +
  ## TTAA
  geom_step(data = ttaa.freq.dt, aes(x = mid, y = TTAA*aK*5),
            size = 0.3, colour = "black") +
  ## Genes
  geom_segment(data = genes.in.tile.dt,
               aes(x = start, xend = end, y=y, yend = y),
               size = 4, colour = "grey50") +
  ## organization
  facet_wrap(~group, nrow = 15) +
  scale_fill_manual(values = group_cols) +
  scale_colour_manual(values = group_cols) +
  scale_y_continuous(limits = c(0,10)) +
  scale_x_continuous(breaks = seq(128200001,132100000,200000)) +
  theme(aspect.ratio = 0.1, legend.position = "none")

## Plot
pcA
```

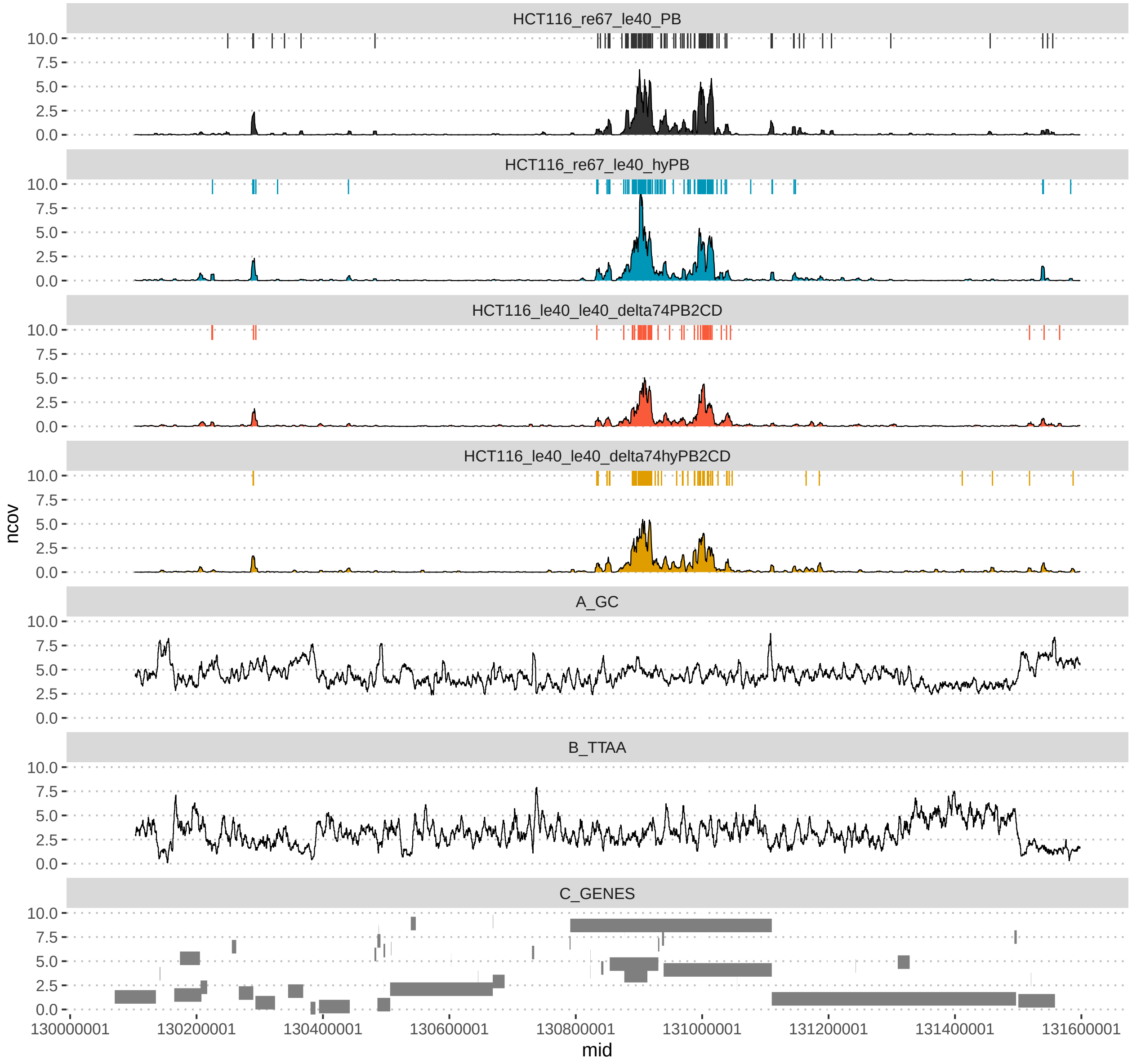


### FUNCTIONS

### filterBamPE()

```
filterBamPE <- function(
                ## INPUTS
                BAM.FILE=NULL,
                BAM.NAME=NULL,
```

```
            STRAND.MODE=2,
            BS.SPECIES=NULL,

            ## FILTERS
            STANDARD.CONTIGS.ONLY=TRUE,
            LIBRARY.MAX.WIDTH=1000,
            QUANTILE.WIDTH.FILTER=TRUE,
            WIDTH.QUANTILE.PROBS=0.999,

            ## OPTIONS
            TAGS=c("NH","NM"),
            SIMPLE.CIGAR=TRUE,
            IS.PAIRED = TRUE,
            IS.DUPLICATE = FALSE,
            IS.PROPER.PAIR = TRUE,
            IS.UNMAPPED.QUERY = FALSE,
            IS.SECONDARY.ALIGNMENT = FALSE,

            ## OPTIONS 2
            YIELD.SIZE=NA
){

## AUTHOR: Pavol Genzor
## Use: Load PE bam file into R
## 10.25.21; Version 3; adding tags, correcting the lane info

## NOTE ON BAM INFO
## 10.25.21; Version 2; added Yield.size and width filter
## 10.22.21; Version 1; original
##

## libraries
suppressPackageStartupMessages({library("data.table");library("dplyr");
  library("Rsamtools");library("GenomicAlignments");
  library("BSgenome.Hsapiens.UCSC.hg38");
  library("BSgenome.Dmelanogaster.UCSC.dm6");
  library("BSgenome.Mmusculus.UCSC.mm10")})

## check input
if(is.null(BAM.FILE)) stop("Please provide full path to a .bam file !!!")
if(is.null(BAM.NAME)) stop("Please provide .bam name !!!")
if(is.null(BS.SPECIES)) stop("Please provide BSSPECIES name !!!")

message("Starting")
message(paste0(" file: ",BAM.NAME))
PROGRESS.L <- list()
PROGRESS.L[["ID"]] <- BAM.NAME

if(isTRUE(STANDARD.CONTIGS.ONLY)){
  WHICH <- keepStandardChromosomes(seqinfo(eval(parse(text = BS.SPECIES))))
} else { WHICH = ""}

message("\tloading parameters")
```

```
message(paste0(
  "\t\t TAGS: ", paste(TAGS,collapse = ", "),"\n",
  "\t\t STRAND.MODE: ",STRAND.MODE,"\n",
  "\t\t SIMPLE.CIGAR: ",SIMPLE.CIGAR,"\n",
  "\t\t IS.PAIRED: ",IS.PAIRED,"\n",
  "\t\t IS.PROPER.PAIR: ",IS.PROPER.PAIR,"\n",
  "\t\t IS.DUPLICATE: ",IS.DUPLICATE,"\n",
  "\t\t IS.SECONDARY.ALIGNMENT: ",IS.SECONDARY.ALIGNMENT,"\n"))

PARAM <- ScanBamParam(which = WHICH,
                      simpleCigar = SIMPLE.CIGAR,
                      tag = TAGS,
                      flag = scanBamFlag(isPaired = IS.PAIRED,
                                         isDuplicate = IS.DUPLICATE,
                                         isProperPair = IS.PROPER.PAIR,
                                         isUnmappedQuery = IS.UNMAPPED.QUERY,
                                         isSecondaryAlignment = IS.SECONDARY.ALIGNMENT))

message("\tloading PE .bam file into GAlignments")
if(is.na(YIELD.SIZE)){message("\t\t all reads")}
else{message(paste0("\t\t YIELD.SIZE: ",YIELD.SIZE))}

PGR <- readGAlignmentPairs(file = BamFile(file = BAM.FILE,
                                          yieldSize = YIELD.SIZE),
                           use.names = TRUE,
                           strandMode = STRAND.MODE,
                           param = PARAM)
PROGRESS.L[["INPUT"]] <- length(PGR)

message("\textracting First read")
FIRST.DT <- as.data.table(first(PGR))
FIRST.DT[,"TAG":=paste("W",width,"NH",NH,"NM",NM,sep = ":")]

message("\textracting Second read")
SECOND.DT <- as.data.table(second(PGR))
SECOND.DT[,"TAG":=paste("W",width,"NH",NH,"NM",NM,sep = ":")]

message("\tcombining First_Second")
FIRST.SECOND <- paste(FIRST.DT[["TAG"]],SECOND.DT[["TAG"]],sep=":")

message("\tmaking GR")
PGR.GR <- granges(PGR)
in.gr <- length(PGR.GR)

message("\tadding tag info")
mcols(PGR.GR)[["TAG"]] <- FIRST.SECOND

if(!is.na(LIBRARY.MAX.WIDTH)){
  message("\tfiltering by width")
  message("\t\tmax width: ",LIBRARY.MAX.WIDTH)
  PGR.GR <- PGR.GR[width(PGR.GR) <= LIBRARY.MAX.WIDTH]
  PROGRESS.L[["LIB_MAX_WIDTH"]] <- length(PGR.GR)}
```

```r
if(isTRUE(QUANTILE.WIDTH.FILTER)){
  message("\tfiltering by quantile")
  message(paste0("\t\tquantile: ",max(WIDTH.QUANTILE.PROBS)))
  WIDTH.QUANTILE <- quantile(x = width(PGR.GR), probs = WIDTH.QUANTILE.PROBS)
  message(paste0("\t\tquantile width: ",max(WIDTH.QUANTILE)))
  PGR.GR <- PGR.GR[width(PGR.GR) <= max(WIDTH.QUANTILE)]
  PROGRESS.L[["QUANTILE_WIDTH"]] <- length(PGR.GR)}

else { message("\tno quantile filter") }

message("\tconverting to DT")
PGR.DT <- as.data.table(PGR.GR)
PGR.DT[,"new_name" := paste(seqnames,start,end,strand,sep = ":")]
PGR.DT <- setDT(PGR.DT)

message("\tinterpreting TAGs")
PGR.DT[,"W1":= tstrsplit(TAG,split=":",fixed=TRUE, keep = 2, type.convert = TRUE)]
PGR.DT[,"W2":= tstrsplit(TAG,split=":",fixed=TRUE, keep = 8, type.convert = TRUE)]
PGR.DT[,"NH1":= tstrsplit(TAG,split=":",fixed=TRUE, keep = 4, type.convert = TRUE)]
PGR.DT[,"NH2":= tstrsplit(TAG,split=":",fixed=TRUE, keep = 10, type.convert = TRUE)]
PGR.DT[,"NM1":= tstrsplit(TAG,split=":",fixed=TRUE, keep = 6, type.convert = TRUE)]
PGR.DT[,"NM2":= tstrsplit(TAG,split=":",fixed=TRUE, keep = 12, type.convert = TRUE)]

## NOTE: This is not a real multiplicity - it tells if two fragments occupy same location
message("\tmaking new DT")
PGR.DT.MULT <- PGR.DT[,.N,by="new_name"]
PGR.DT.MULT[,"chr" := tstrsplit(new_name,split=":",keep = 1)]
PGR.DT.MULT[,"start" := tstrsplit(new_name,split=":",keep = 2)]
PGR.DT.MULT[,"end" := tstrsplit(new_name,split=":",keep = 3)]
PGR.DT.MULT[,"strand" := tstrsplit(new_name,split=":",keep = 4)]
colnames(PGR.DT.MULT) <- gsub("N","MULT",colnames(PGR.DT.MULT))

message("\tsummarizing TAGs")
PGR.DT.W <- PGR.DT[,lapply(.SD,min),by="new_name",.SDcols = c("W1","W2")]
PGR.DT.NH <- PGR.DT[,lapply(.SD,max),by="new_name",.SDcols = c("NH1","NH2")]
PGR.DT.NM <- PGR.DT[,lapply(.SD,function(s){sum(s)/length(s)}),by="new_name",
                    .SDcols = c("NM1","NM2")]

message("\tcombining DTs")
PGR.DT.M.W <- PGR.DT.MULT[PGR.DT.W,on=c("new_name")]
PGR.DT.M.W.NH <- PGR.DT.M.W[PGR.DT.NH,on=c("new_name")]
PGR.DT.M.W.NH.NM <- PGR.DT.M.W.NH[PGR.DT.NM,on=c("new_name")]
PGR.DT.M.W.NH.NM[["new_name"]] <- NULL

message("\tmaking new GR")
PGR.NGR <- makeGRangesFromDataFrame(df = PGR.DT.M.W.NH.NM, keep.extra.columns = TRUE)

message("\treporting")
PROGRESS.L[["FINAL UNIQUE FRAGMENTS"]] <- length(PGR.NGR)
PROGRESS.DT <- melt.data.table(as.data.table(PROGRESS.L), id.vars = "ID",
                               value.name = "count", variable.name = "step")
PROGRESS.DT[["percent"]] <- (PROGRESS.DT[["count"]]/
                               PROGRESS.DT[step %in% "INPUT"][["count"]])*100
```

```
  write.csv(file=paste0(getwd(),"/",BAM.NAME,"_filterBamPE.tab"),x = PROGRESS.DT,
            quote = FALSE, row.names = FALSE)

  message("Done!")
  message("")
  print(PROGRESS.DT)
  return(PGR.NGR)
}
```

**findPeaksInPERegions()**

```
findPeaksInPERegions <- function(IN.GR=NULL,
                                 GR.NAME=NULL,
                                 MIN.READS=5,
                                 PPM.FACTOR=1000000){

  ## GOAL: find simple stranded peaks in Genomic Ranges
  ## Author: Pavol Genzor
  ## 04.20.22; Version 1

  ## Libraries
  suppressPackageStartupMessages({library(data.table); library(IRanges);
    library(GenomicRanges); library(GenomicAlignments)})

  ## Input checking
  if(is.null(IN.GR)) stop("Please provide IN.GR!")
  if(is.null(GR.NAME)) stop("Please provide GR.NAME!")
  if(!"NH1" %in% colnames(mcols(IN.GR)) | !"NH2" %in% colnames(mcols(IN.GR)))
    stop("This is not PE data. You need NH1 and NH2 columns!")

  message("Looking in sample:")
  message(paste0("  ",GR.NAME))

  message("\tusing unique anchored")
  NGR <- IN.GR[mcols(IN.GR)[["NH1"]] %in% 1 | mcols(IN.GR)[["NH2"]] %in% 1]
  total_regions <- length(NGR)

  message("\tcalculating stranded coverage")
  COV.PLUS <- GenomicRanges::coverage(NGR[strand(NGR) %in% "+"])
  COV.MINUS <- GenomicRanges::coverage(NGR[strand(NGR) %in% "-"])

  message("\tfinding peaks")
  message(paste0("\t\tmin. read: ", MIN.READS))
  SLICE.PLUS <- IRanges::slice(x = COV.PLUS, lower = MIN.READS)
  SLICE.MINUS <- IRanges::slice(x = COV.MINUS, lower = MIN.READS)

  message("\tmaking range with strand")
  PLUS.GR <- GenomicRanges::reduce(as(SLICE.PLUS, "GRanges"))
  MINUS.GR <- GenomicRanges::reduce(as(SLICE.MINUS, "GRanges"))
  strand(PLUS.GR) <- "+";  strand(MINUS.GR) <- "-"

  # adding total sequences
```

```r
  mcols(PLUS.GR)[["total_regions"]] <- total_regions
  mcols(MINUS.GR)[["total_regions"]] <- total_regions

  message("\tcounting sequences")
  mcols(PLUS.GR)[["uniq_region_count"]] <- GenomicRanges::countOverlaps(
    query = PLUS.GR, subject = NGR)
  mcols(MINUS.GR)[["uniq_region_count"]] <- GenomicRanges::countOverlaps(
    query = MINUS.GR, subject = NGR)

  message("\tadding peak info")
  message(paste0("\t\tmaximum peak height"))
  mcols(PLUS.GR)[["peak_height"]] <- unlist(rbindlist(lapply(
    SLICE.PLUS,function(x){as.data.table(max(x))})))
  mcols(MINUS.GR)[["peak_height"]] <- unlist(rbindlist(lapply(
    SLICE.MINUS,function(x){as.data.table(max(x))})))

  message(paste0("\t\traw area under the curve"))
  mcols(PLUS.GR)[["peak_auc"]] <- unlist(rbindlist(lapply(
    SLICE.PLUS,function(x){as.data.table(sum(x))})))
  mcols(MINUS.GR)[["peak_auc"]] <- unlist(rbindlist(lapply(
    SLICE.MINUS,function(x){as.data.table(sum(x))})))

  message(paste0("\t\tnormalized area under the curve"))
  mcols(PLUS.GR)[["peak_nauc"]] <- as.numeric(
    mcols(PLUS.GR)[["peak_auc"]] / (total_regions/PPM.FACTOR))
  mcols(MINUS.GR)[["peak_nauc"]] <- as.numeric(
    mcols(MINUS.GR)[["peak_auc"]] / (total_regions/PPM.FACTOR))

  message(paste0("\t\tpeak width"))
  mcols(PLUS.GR)[["peak_width"]] <- width(PLUS.GR)
  mcols(PLUS.GR)[["sample"]] <- GR.NAME
  mcols(MINUS.GR)[["peak_width"]] <- width(MINUS.GR)
  mcols(MINUS.GR)[["sample"]] <- GR.NAME

  message("Finished.")
  aPEAKGR <- c(PLUS.GR,MINUS.GR)
  aPEAKGR <- sort.GenomicRanges(aPEAKGR)
  return(aPEAKGR)
}
```
