## Supplementary Data for "Transposase N-terminal phosphorylation and asymmetric transposon ends inhibit *piggyBac* transposition in mammalian cells"

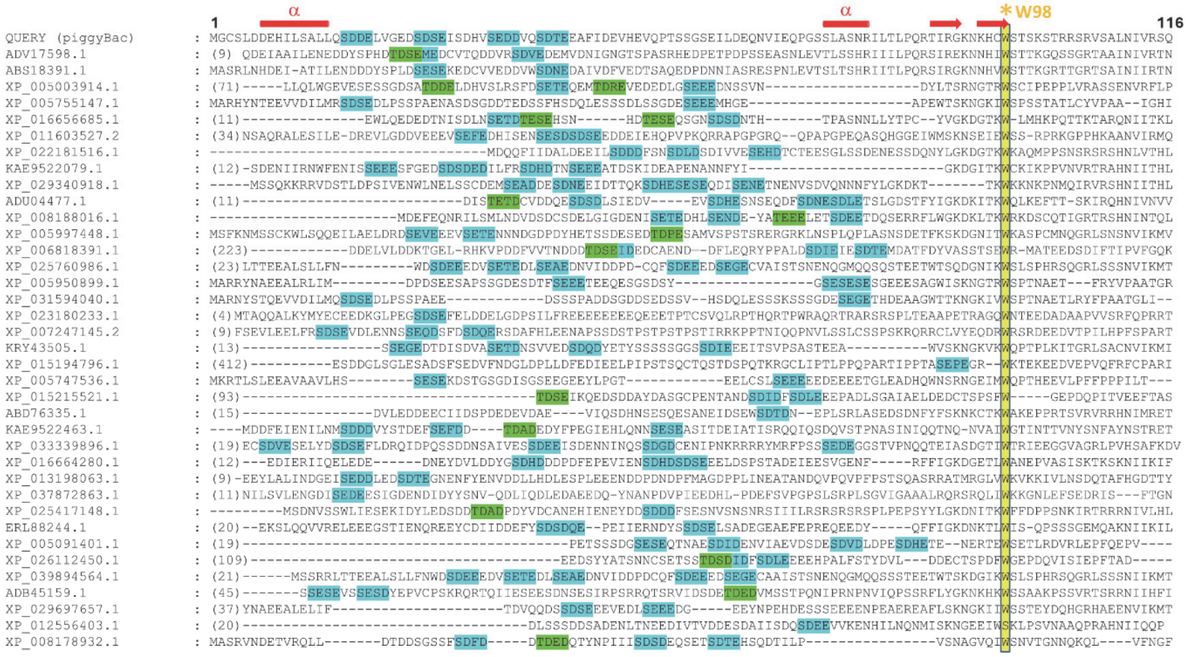


**SFig. 1 N-termini of PBLEs can contain multiple CKII phosphorylation motifs**

The sequence alignment was generated by JPred4 using the PB transposase N-terminus as a query, and manually adjusted in the region of AA W98. Genbank or NCBI reference numbers are shown on the left. Sequences that match the canonical CKII motif (pS/pT-D/E-X-D/E) are highlighted in blue or green. The secondary structure prediction for PB (amino acids numbered) by AlphaFold2 is shown schematically across the top in red.

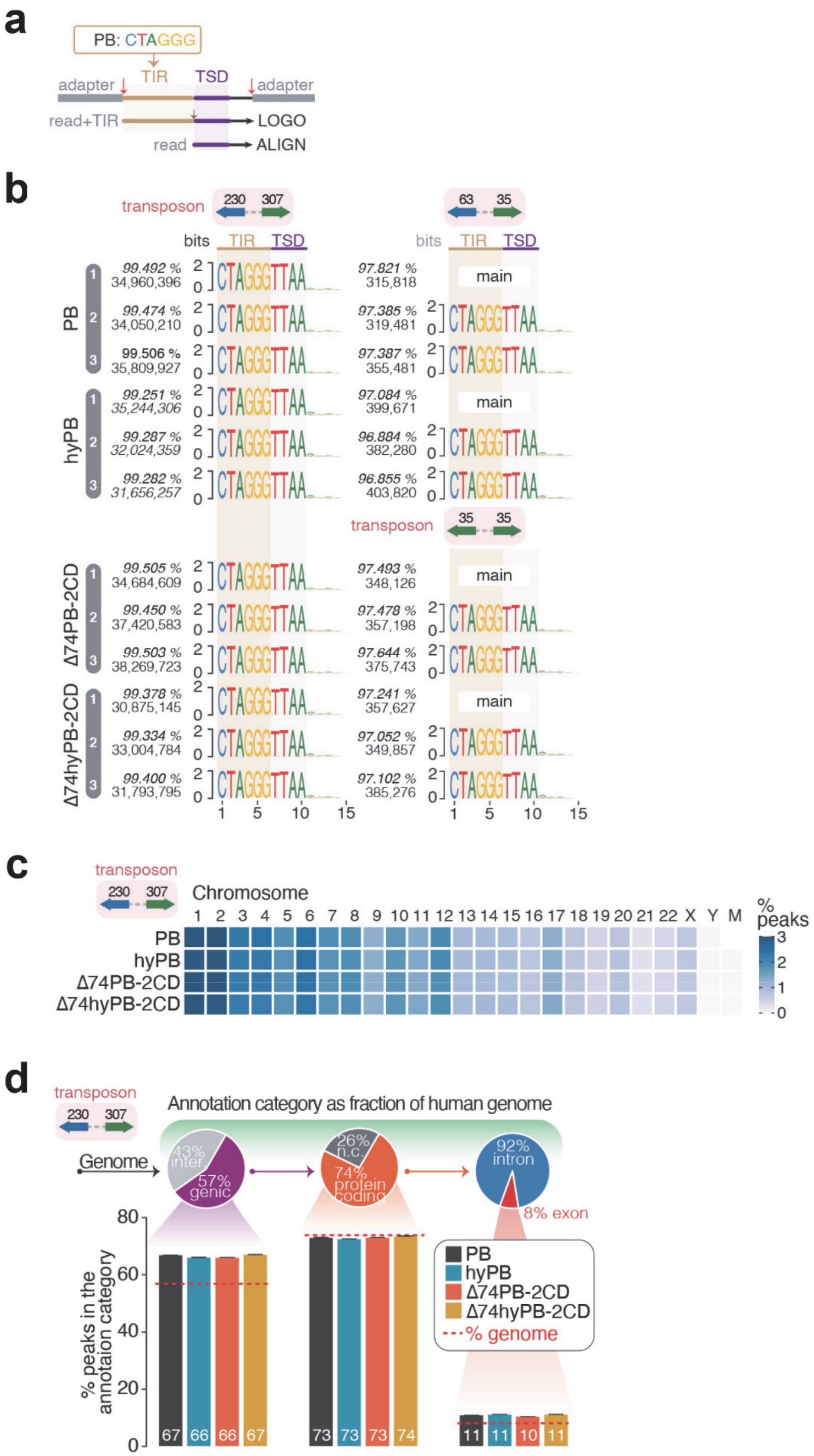
a
PB: CTAGGG
adapter
TIR
TSD
adapter
read+TIR
LOGO
read
ALIGN
b
transposon
230
307
63
35
bits
TIR
TSD
PB
hyPB
Δ74PB-2CD
Δ74hyPB-2CD
99.492 % 34,960,396
99.474 % 34,050,210
99.506 % 35,809,927
99.251 % 35,244,306
99.287 % 32,024,359
99.282 % 31,656,257
99.505 % 34,684,609
99.450 % 37,420,583
99.503 % 38,269,723
99.378 % 30,875,145
99.334 % 33,004,784
99.400 % 31,793,795
97.821 % 315,818
97.385 % 319,481
97.387 % 355,481
97.084 % 399,671
96.884 % 382,280
96.855 % 403,820
97.493 % 348,126
97.478 % 357,198
97.644 % 375,743
97.241 % 357,627
97.052 % 349,857
97.102 % 385,276
main
CTAGGGTTAA
1 5 10 15
c
transposon
230 307
Chromosome
1 2 3 4 5 6 7 8 9 10 11 12 13 14 15 16 17 18 19 20 21 22 X Y M
PB
hyPB
Δ74PB-2CD
Δ74hyPB-2CD
% peaks
3 2 1 0
d
transposon
230 307
Annotation category as fraction of human genome
Genome
43% inter
57% genic
26% n.c.
74% protein coding
92% intron
8% exon
% peaks in the annotaion category
0 20 40 60 80
67 66 66 67
73 73 73 74
11 11 10 11
PB
hyPB
Δ74PB-2CD
Δ74hyPB-2CD
% genome

**SFig. 2 Genome-wide characterization of re-engineered transpososomes.**
**a.** A schematic representation of the initial computational processing of re-engineered transpososome reads. First, the machine adapters were trimmed and the reads containing TIR sequences were used to generate sequence logos. Next, TIR sequence was removed and reads aligned to genome. **b.** Excision profiles and target site duplication (TSD) remains unchanged between wild-type and mutant transpososome systems. Sequence logos for the first 15 nucleotides of the reads with TIR sequence are shown. The TSD (TTAA) is indicated. Percentage of all reads with perfect TIR and final read number in analysis are shown on the left. **c.** Genome-wide distribution of peaks across different chromosomes remained unchanged between mutant proteins when evaluated using transposons system with long LE and RE arms. The fraction of insertion peaks is shown as heatmap for individual chromosomes. **d.** Annotation of peaks overlapping common genome annotation categories within the human genome (Gencode annotation version 24) in experiments with long LE and RE arms indicate conservation of integration site preferences.

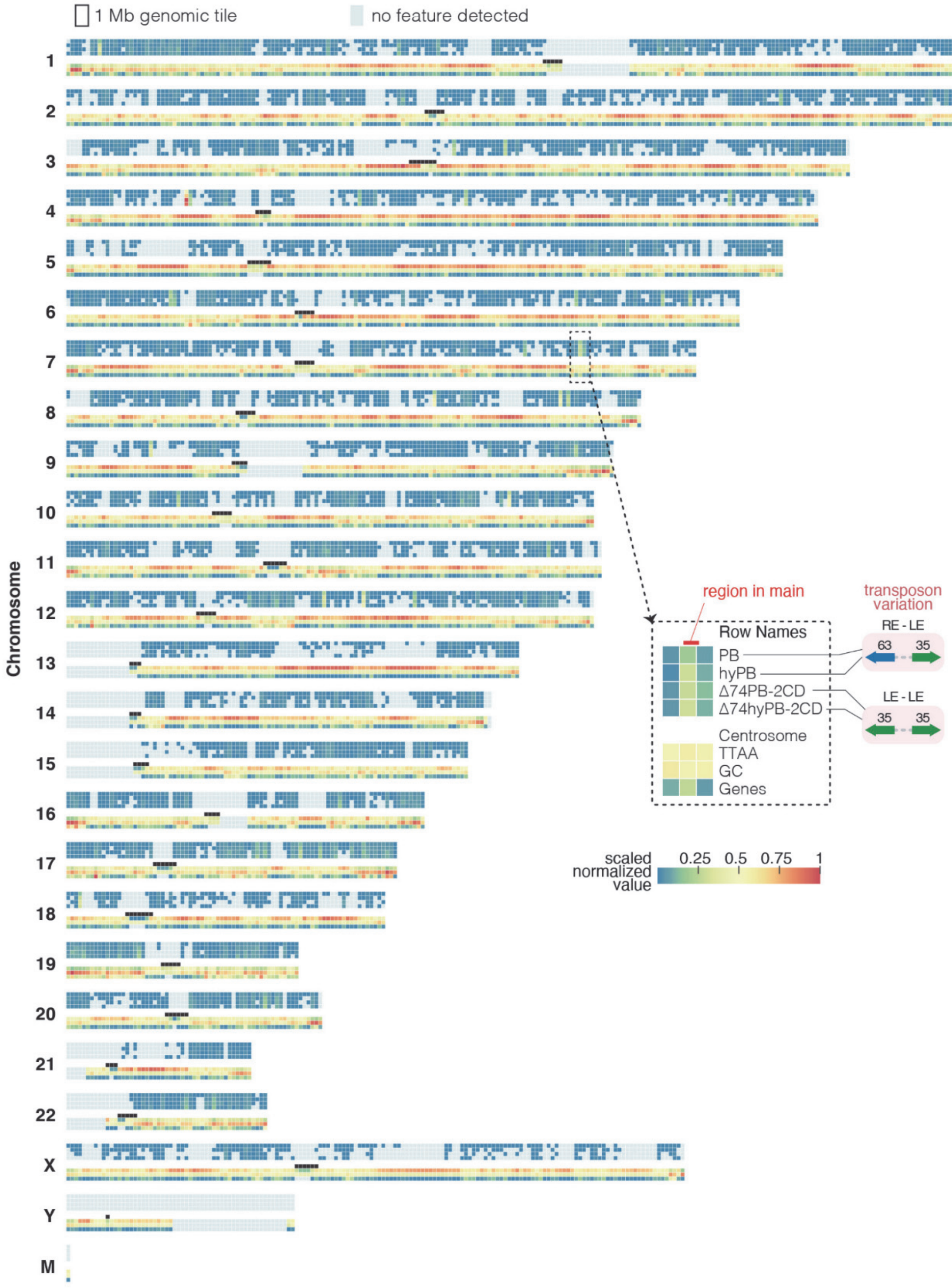


**SFig. 3 Genome-wide map of PB integrations in the human genome**
A one mega-base tile map of entire human genome showing normalized abundance of the PB and peaks, and genomic features including location of Centrosomes, normalized TTAA and GC

frequencies, and normalized gene abundance. Insert is highlighting an approximate location of region described in main figure.

**SData. Excel spreadsheet of identified phosphorylated PB peptides**

If the Modscore values for Site 1 Position_A and Site 1 Position B are both above 19 for the same site (highlighted in pink) then the location is considered confidently assigned. In the case of multiple phosphorylation events the Site 1 Position refers to the N-terminal most site and Site 2 Position refers to the next site moving toward the C-terminal. Green highlights indicate lowered confidence in peptide assignment.

**Supplementary table:** Primers for various analyses, sequences 5'-to-3'

| | |
|---|---|
| **Excision assay** | |
| ExcF | ATGCGGCATCAGAGCAGATT |
| ExcR | TGTGTGGAATTGTGAGCGGA |
| **Plasmid rescue** | |
| LEseq | TAAGCCCACTGCAAGCTACC |
| REseq | CCTCCAAGCCAGTTACCTCG |
| **qPCR copy number analysis** | |
| NeoF | CGGATGGAAGCCGGTCTTGTC |
| NeoR | TCATTTTGACTCACGCGG |
| Neo probe | GCGTTGGCTACCCGTGATATTGC 5' 6-FAM/ZEN/3' IB®FQ |
| RNasePF | AGATTTGGACCTGCGAGCG |
| RNasePR | AGAAGGCGATAGAAGGCGATG |
| RNaseP probe | TTCTGACCTGAAGGCTCTGCGCG 5'-6-HEX /ZEN/ 3' IB®FQ |
| **Sequencing library preparation** | |
| SMART-dT18VN | AAGCAGTGGTATCAACGCAGAGTACGTTTTTTTTTTTTTTTTTTTTTTTTTTTTTVN |
| SRT-PAC-F1 | CAACCTCCCCTTCTACGAGC |
| SRT-Seq P1 | CGGTGCCTGATAACAACTCG |
| SRT-Seq P2 | GGTTCAGCAGGAATGCCGAG |
| SRT-Seq P3 | CTCAGTGTCCATAGCGACGC |
| Smart | AAGCAGTGGTATCAACGCAGAGT |
| Read1-TnME | ACACTCTTTCCCTACACGACGCTCTTCCGATCTAGATGTGTATAAGAGACA*G |
| Read2-R-5'TIR | GACTGGAGTTCAGACGTGTGCTCTTCCGATCTTTTACGCAGACTATCTTTCTAG |
